# Dynamic Phosphorylation of RGS Provides Spatial Regulation of MAP Kinase and Promotes Completion of Cytokinesis during the Yeast Pheromone Response

**DOI:** 10.1101/2021.06.22.449324

**Authors:** William C. Simke, Cory P. Johnson, Andrew J. Hart, Sari Mayhue, P. Lucas Craig, Savannah Sojka, Joshua B. Kelley

**Author notes:** Corresponding Author: Joshua Kelley. These authors contributed equally (William C. Simke, Cory P. Johnson).

## Abstract

Yeast use a G-protein coupled receptor (GPCR) signaling pathway to detect mating pheromone, arrest in G1, and direct polarized growth towards the potential mating partner. The primary negative regulator of this pathway is the regulator of G-protein signaling (RGS), Sst2, which induces Gα GTPase activity and subsequent inactivation of all downstream signaling including a MAPK cascade. The MAPK Fus3 phosphorylates the RGS in response to pheromone, but the role of this modification is unknown. We set out to examine the role of RGS phosphorylation during the pheromone response. We found that RGS phosphorylation peaks early in the pheromone response and diminishes RGS localization to the polarity site and focuses MAPK complexes there. At later time points, RGS is predominantly unphosphorylated, which promotes RGS localization to the polar cap and broadens the distribution of MAPK complexes relative to the Cdc42 polarity machinery. Surprisingly, we found that phosphorylation of the RGS is required for the completion of cytokinesis prior to pheromone induced growth. The completion of cytokinesis in the presence of pheromone is promoted by the formin Bnr1 and the kelch-repeat protein, Kel1, both proteins previously found to interact with the RGS.

## Introduction

When cells receive signals to perform competing processes, they must integrate those two signals into a singular outcome. The budding yeast, *Saccharoymyces cerevisiae*, use a G-protein coupled receptor (GPCR) to detect and grow toward potential mating partners (Alvaro and Thorner, 2016; Arkowitz, 2009; Wang and Dohlman, 2004). However, the cells must complete mitosis and arrest in G1 prior to mating projection morphogenesis (shmoo formation) (Peter et al., 1993). This requires that the cell prioritize the signaling that will drive mitosis and cytokinesis to completion, and only after arrest in G1, allow the pheromone signaling pathway to commandeer the Cdc42 polarity machinery that has shared uses in both mitosis and pheromone-induced morphogenesis (Chiou et al., 2017; Park and Bi, 2007). While the mechanism by which G1 arrest occurs is understood, the mechanism responsible for suppression of receptor driven polarization is unknown.

The pheromone response can be thought of as a response made up primarily of two G-proteins: the receptor-activated large G-protein consisting of the Gα and Gβγ conveying information about where the pheromone receptor is active, and the small G-protein Cdc42 controlling MAPK signaling and actin cytoskeleton polarization. The GPCR Ste2 activates the large G-protein, causing the Gα and Gβγ subunits to dissociate. Gβγ initiates Cdc42 mediated polarization of the actin cytoskeleton to form a mating projection. Gβγ also promotes the activation of the two yeast ERK homologs, Fus3 and Kss1(Wang and Dohlman, 2004). Of these two MAP kinases, Fus3 has pheromone specific roles: it is required for gradient tracking, arrest of the cell cycle in G1, and is scaffolded to the cell periphery by active Gα to regulate actin polymerization (Elion et al., 1993; Hao et al., 2008; Matheos et al., 2004; Metodiev et al., 2002; Pope et al., 2014). For this study we will only be concerned with Fus3 functions, and so all references to MAPK refer to Fus3.

The primary negative regulator of the pheromone pathway is the Regulator of G-protein Signaling (RGS), Sst2, (Chasse et al., 2006) which serves as the GTPase activating protein (GAP) for the Gα subunit (Apanovitch et al., 1998). Upon hydrolyzing GTP, the Gα binds to Gβγ, turning off the pathway. RGS function is required for pathway inactivation and for the ability of the cell to track the pheromone gradient (Dohlman et al., 1995; Segall, 1993). GPCR signaling pathways play a central role in human disease, indicating that understanding the mechanisms by which RGS functions has the potential to inform understanding of human signaling pathways relevant for drug development (Bar-Shavit et al., 2016; Hauser et al., 2017; Lappano and Maggiolini, 2012).

The RGS Sst2 has characterized interactions with the Gα (Gpa1) subunit, the pheromone receptor (Ste2), and the MAPK (Fus3) (Ballon et al., 2006; DiBello et al., 1998; Garrison et al., 1999). RGS serves as a GTPase activating protein for Gα, a function that is enhanced by its binding to the cytoplasmic tail of the receptor (Apanovitch et al., 1998; Ballon et al., 2006; Dixit et al., 2014). The MAPK Fus3 phosphorylates the RGS at serine 539 in a pheromone dependent manner, but this does not impact the sensitivity of the pathway or downstream MAPK or transcriptional outputs (Garrison et al., 1999).

In less well-characterized interactions, the RGS has been found in a yeast two-hybrid screen to interact with the formin Bnr1 and the formin regulatory protein Kel1 (Burchett et al., 2002; Yu et al., 2008). Yeast contain two formins, Bni1 and Bnr1, which drive polymerization of actin in response to Rho family signaling (Breitsprecher and Goode, 2013). Bni1 is activated by Cdc42, and is known to be involved in the pheromone response as its activity is regulated by MAPK bound to active Gα (Evangelista et al., 1997; Matheos et al., 2004; Metodiev et al., 2002). Bnr1 is activated by Rho3 and Rho4, and is involved in cytokinesis, but has no clear role in the pheromone response (Imamura et al., 1997). The formins are partially redundant, as either is sufficient for viability, but the loss of both is lethal (Imamura et al., 1997). Kel1 is a kelch-repeat containing protein which has been shown to act as a negative regulator of Bnr1, is required for efficient mating, and plays a role in the mitotic exit network (Gould et al., 2014; Hofken and Schiebel, 2002; Smith and Rose, 2016). Given the clear roles of Bnr1 and Kel1 in cytokinesis with little known about their roles in the pheromone response, the potential for RGS interactions with these proteins has not been pursued.

In an unsynchronized, logarithmically growing population of yeast, most cells will be between the beginning of S phase and cytokinesis. Because the Cdc42 polarity machinery is shared between both the mitotic machinery and the pheromone machinery, cells must arrest in G1 before responding to pheromone (Bi and Park, 2012; Chiou et al., 2017). This is accomplished by MAPK phosphorylating Far1, which inhibits Cyclin Dependent Kinase (Butty et al., 1998; Peter et al., 1993; Pope et al., 2014). Thus, when cells receive the pheromone signal, they complete their current round of mitosis and cytokinesis, and only then does receptor driven signaling reorient the polarity machinery for pheromone-induced morphogenesis of the mating projection (shmoo) (Chiou et al., 2017).

Here, we set out to determine the role of MAPK phosphorylation of RGS in response to pheromone. We found that RGS phosphorylation is dynamic, with high phosphorylation early in the response, followed by decreased phosphorylation later. Phosphorylation of the RGS decreases its localization to the polar cap, and reduces the distance between peak active Cdc42 and peak MAP Kinase localization. Surprisingly, RGS phosphorylation early in the pheromone response is required to ensure that cytokinesis completes prior to the beginning of pheromone induced polarity. We find that cytokinetic completion in the presence of pheromone is also dependent upon the kelch-repeat protein Kel1, a component of the mitotic exit network.

## Results

### Cells can track a gradient of pheromone independent of phosphorylation at Ser 539

The ability to track a gradient of pheromone is dependent upon RGS, specifically its GTPase activating protein (GAP) function (Dixit et al., 2014; Segall, 1993). MAPK phosphorylates RGS on serine 539 in response to pheromone, though the nature of this phosphorylation event is poorly understood. While previous studies found that GAP activity was not affected by S539 phosphorylation, it is possible that phosphorylation might change Sst2 function in time or space in a way that affects gradient tracking (Garrison et al., 1999). We tested this hypothesis by using strains expressing an unphosphorylatable RGS mutant (*sst2^S539A^*, denoted RGS) and phospho-mimetic RGS mutant (*sst2^S539D^*, denoted p*RGS) from the endogenous SST2 genomic locus, each fused to GFP. As a marker of the polar cap, we used the Cdc42-GTP binding protein Bem1 fused to mRuby2 (Kelley et al., 2015). These strains were examined in a microfluidic gradient chamber by live cell microscopy (Suzuki et al., 2021). We exposed these cells to a 0-150 nM gradient of pheromone and measured their ability to grow toward the source of pheromone (Figure 1). Both phospho-mutant strains were able to track a gradient of pheromone (Figure 1B). Thus, feedback phosphorylation of RGS has no effect on gradient tracking.

**Figure 1.**
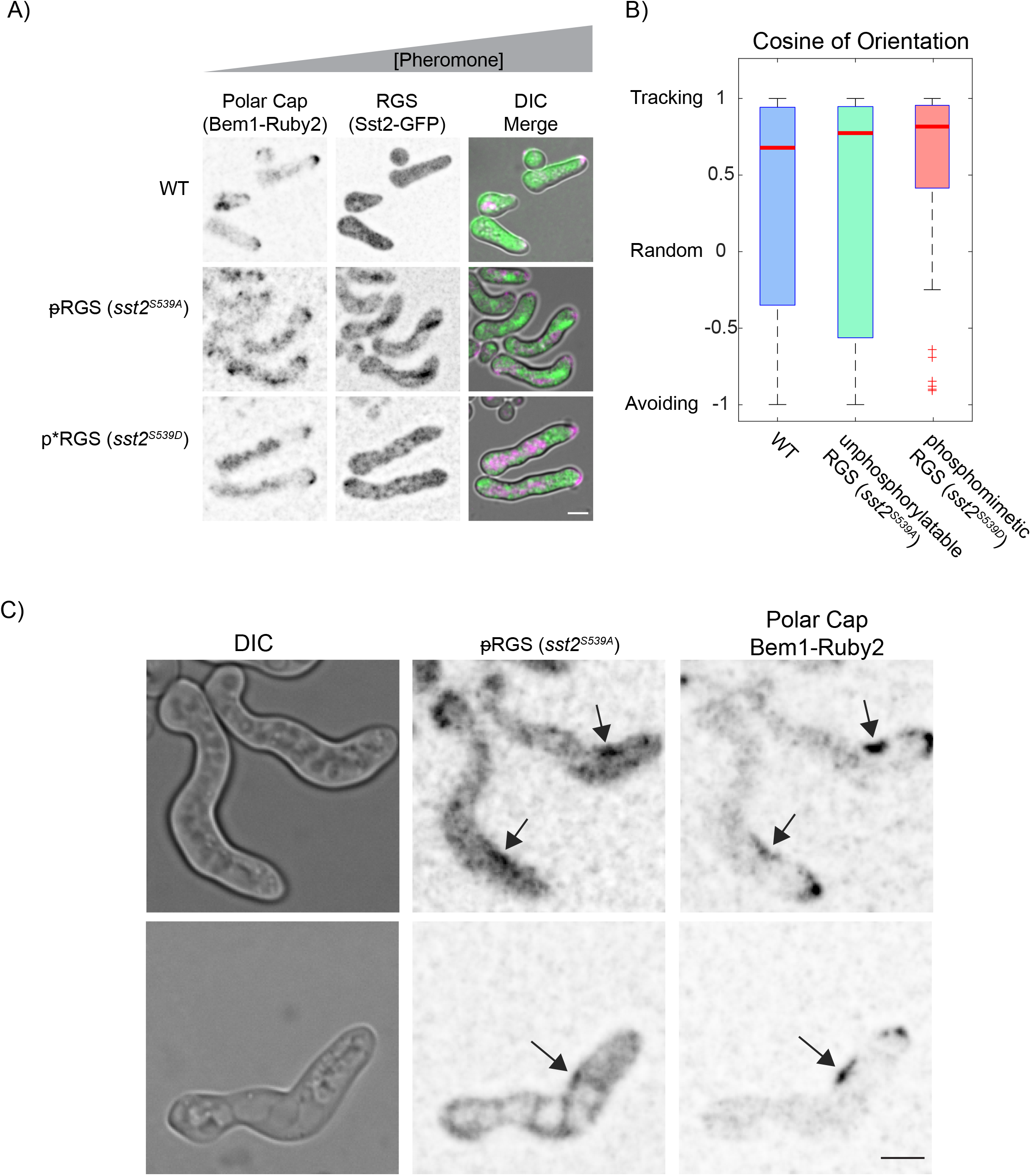
Phosphorylation state of the RGS does not stop gradient tracking. A) Representative live-cell images of WT, unphosphorylatable (RGS), and phosphomimetic (p*RGS) RGS expressing the polar cap marker (Bem1-mRuby2) and the RGS (Sst2-EGFP) tracking a 0-150nM gradient of pheromone, with pheromone increasing to the right. B) Quantification of gradient tracking cells measured by the cosine of orientation for WT (n =95), RGS (*sst2^S539A^*, n =39), and p*RGS (*sst2^S539D^*, n =45) from three experiments. Scale bars represent 5 µm. C) Examples of the polar cap marker Bem1 colocalizing with the RGS at sites distal to the polar cap in the unphosphorylatable RGS mutant *sst2^S539A^*.

In the unphosphorylatable RGS mutant (*sst2^S539A^*), the polar cap occasionally colocalized with the RGS at the inside of turns (Figure 1B). Normally, septins and RGS accumulate at the inside of turns (Kelley et al., 2015), but the polar cap does not. This suggests a defect in restricting the polar cap in the unphosphorylatable mutant.

### RGS localization is regulated by its phosphorylation state

The pheromone driven MAPK Fus3 is central to mating signaling, driving cell cycle arrest, transcriptional changes, and promoting actin cytoskeleton changes (Alvaro and Thorner, 2016; Chiou et al., 2017; Wang and Dohlman, 2004) (Garrison et al., 1999). We hypothesized that Fus3 phosphorylation of the RGS may regulate its spatial distribution. To test this, we again used the p*RGS (*sst2^S539D^*) and RGS (*sst2^S539A^*) mutants tagged with EGFP, as well as Bem1-mRuby2 for polar cap localization. Cells were exposed to saturating pheromone (300 nM) in a microfluidic chamber (Suzuki et al., 2021) for 12 hours and imaged by timelapse epifluorescence microscopy. To examine the distribution of the RGS along the periphery of the cell, we used our previously reported approach of spatial normalization to the polar cap (Kelley et al., 2015; Shellhammer et al., 2019). Briefly, the signal of RGS along the periphery of the cell is spatially registered to the center of the polar cap as identified by peak Bem1 signal, and then averaged to generate a distribution of the protein during the pheromone response (Figure 2) (Kelley et al., 2015). Fluorescence intensity was normalized to sum to 1, so that the values shown indicate the average fraction of protein found at that position relative to the center of the polar cap. Wild type RGS localizes to the polar cap and to the periphery of projections where septins would be, consistent with our previous findings (Figure 3A, B) (Dixit et al., 2014; Kelley et al., 2015). The amount of RGS at the polar cap varied, appearing to come and go, while its association with the base of the mating projection was more consistent (Figure 3B). We found that the phospho-mimetic mutation diminishes RGS localization to the polar cap (Figure 3A, B). The inability to phosphorylate RGS leads to a small but statistically significant increase in association with the polar cap, evident in the images as a more consistently discernable concentration of RGS at the tip of the shmoo. The similarity between the profiles of WT and unphosphorylatable RGS suggests that much of the RGS measured in WT cells may be in the unphosphorylated form (Figure 3C). We then examined changes in RGS distribution using an averaged 3D-kymograph (Figure 3D). As expected, RGS fluorescence increases throughout the time course, which is due to persistent pheromone-induced production of RGS (Dohlman et al., 1996). As the time in pheromone passes 400 minutes, there is broadening of the lower intensity area surrounding the polar cap in both WT and RGS cells, but a consistent and discernable peak of intensity at the center of the polar cap for both WT and the RGS mutant that is not evident in the p*RGS (also clear in Figure 3B). This further suggests that the phospho-form of the RGS is unable to interact with a normal binding partner at the center of the polar cap.

**Figure 2.**
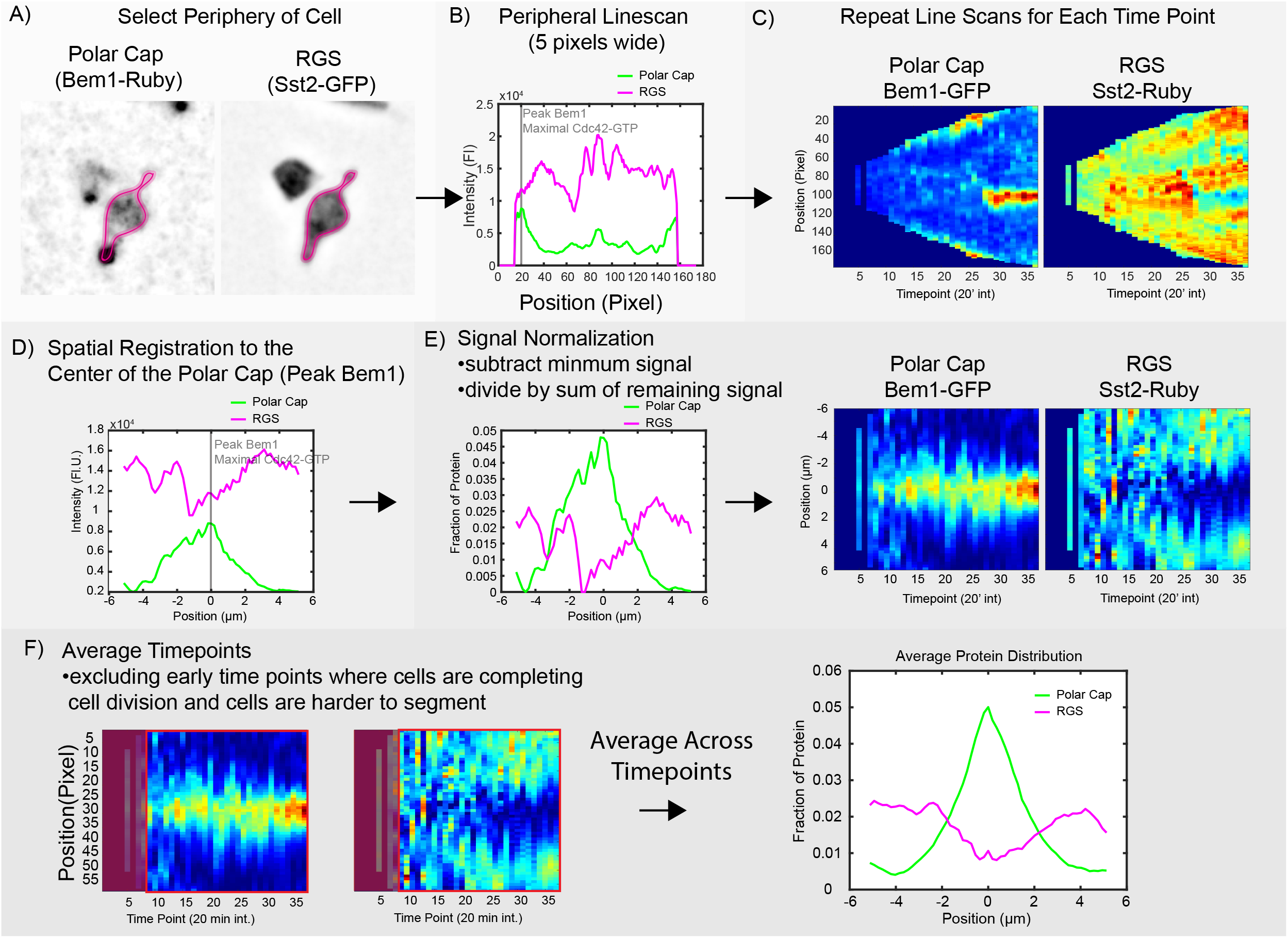
Method for determining protein distribution on the cell periphery. A) The periphery of the cell is defined for a specific time point. B) The intensity of fluorescence signal from each channel is measured with a line width of 5 pixels. C) The periphery for each channel at every time point is measured, here shown as a kymograph. D) The linescans are adjusted such that the peak intensity of the polar cap marker Bem1 is set to the center, and the other channel is moved to maintain its original spatial relationship with bem1. E) The intensity profiles are normalized by subtracting the minimum value and then dividing by the total intensity in the line such that the new line sums to 1. F) For the average distributions, all lines after time point 9 are averaged. Time points before this often still include cells complete cytokinesis, and are less easily segmented.

**Figure 3.**
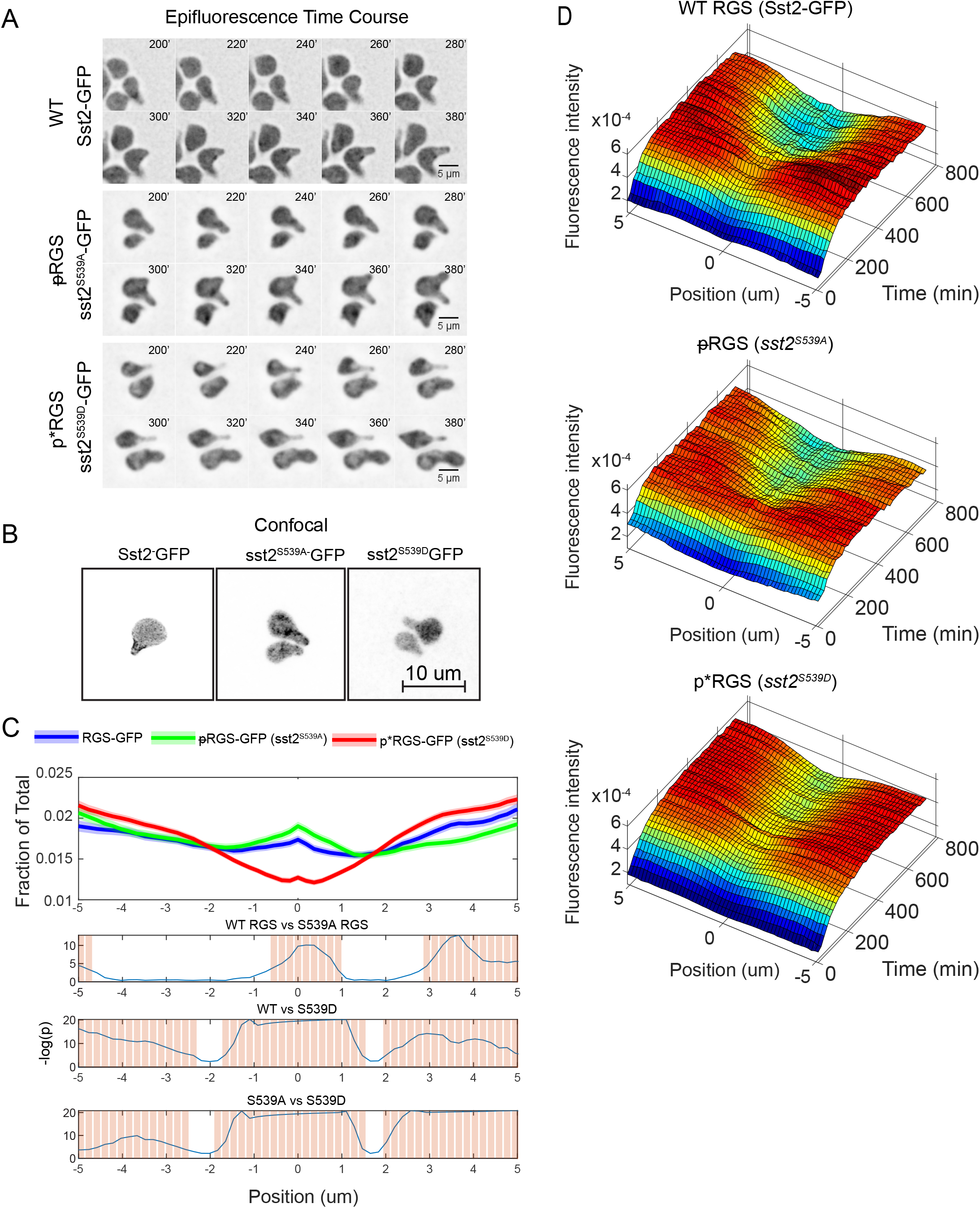
Localization of the RGS is dependent on phosphorylation state. A) Epifluorescence time course images of the strains expressing the indicated RGS mutants (Sst2-GFP) imaged in a microfluidic device exposed to 300 nM pheromone for the indicated time. B) Confocal images of WT, unphosphorylatable (RGS), and phosphomimetic (p*RGS) RGS fused to EGFP in saturating pheromone (10 µM). C) Quantification of the average RGS spatial distribution normalized to the polar cap marker (Bem1-mRuby2) in saturating pheromone over a 12hr time-course in a microfluidic gradient chamber, imaged by epifluorescence microscopy. Lines are derived from averaging from 180 min. onward. Bottom graphs display statistical analysis using one-way ANOVA followed by Tukey’s HSD, with statistically (p < 0.05) significant differences in localization noted by bars. Data is derived from n = 89 cells (WT), n = 88 RGS (unphosporylatable), and n = 139 p*RGS (phosphomimetic) cells per strain, with 29 time points per cell. D) 3-D kymographs of the spatial distribution of the RGS over 12hrs for WT, RGS, and p*RGS with 37 time points per cell from (A).

### RGS phosphorylation alters the distance between the Polar Cap and the Gα-MAPK complex

While much of the polarity signaling in the pheromone response comes through the Gβγ branch of the pathway, GTP bound Gα is known to recruit active MAPK (Metodiev et al., 2002). The Gα-MAPK complex activates the formin Bni1 and promotes gradient tracking (Errede et al., 2015; Matheos et al., 2004; Metodiev et al., 2002). We hypothesized that the phospho-dependent changes in RGS localization would lead to corresponding changes in the localization of active Gα. While Gα is localized across the membrane (Wang et al., 2005), its localization alone does not indicate whether Gα is activated. However, we hypothesized that the activation state of Gα would influence MAPK localization in a measurable way.

To test the use of MAPK as a maker for active Gα, we examined a GFP tagged MAPK (Fus3-GFP) in wild type cells, and two mutants with opposing effects: a Gα mutant that does not bind MAPK, *gpa1^E21 E22^* (Metodiev et al., 2002); and a Gα mutant that is hyperactive because it no longer interacts with RGS, *gpa1^G302S^* (Figure 4A)(DiBello et al., 1998). In WT cells, MAPK localizes to the nucleus and to the polar cap, although much like the RGS, the association with the polar cap fluctuated in intensity (Figure 4B). In the *gpa1^E21E22^* mutant that cannot bind MAPK, there was a marked decrease in association with the polar cap, although at later time points, it appeared at the polar cap more often (Figure 4B, D). In the hyperactive *gpa1^G302S^* MAPK levels consistently increased, and there were often multiple discernable spots of MAPK accumulation (Figure 4B). We assessed MAPK levels along the periphery of the cell in these strains using the same techniques as above, with the exception that the nuclei were masked to exclude nuclear signal from our measurements of the periphery (Figure 4C). Nuclear masks were generated by using single cell histogram analysis and removing large objects that were more than 1 standard deviation above the mean, an adaptation of the algorithm designed to detect granules (Hunn et al., 2021). When examining the shapes of the protein distributions, it is clear that WT cells have the sharpest distribution of MAPK with respect to the location of the polar cap. Both loss of MAPK binding and excess binding of MAPK broadened its distribution (Figure 4E). We believe the distribution of signal in the Gα MAPK-binding mutant (*gpa1^E21E22^*) likely represents the profile of the remaining MAPK binding partners at the polar cap (Figure 4D). Fus3 binds to the MAPK scaffold Ste5, and the polar cap is populated with many MAPK substrates (Elion et al., 1993; Wang and Dohlman, 2004). We conclude that MAPK localization is affected by the state of its binding partner, Gα.

**Figure 4.**
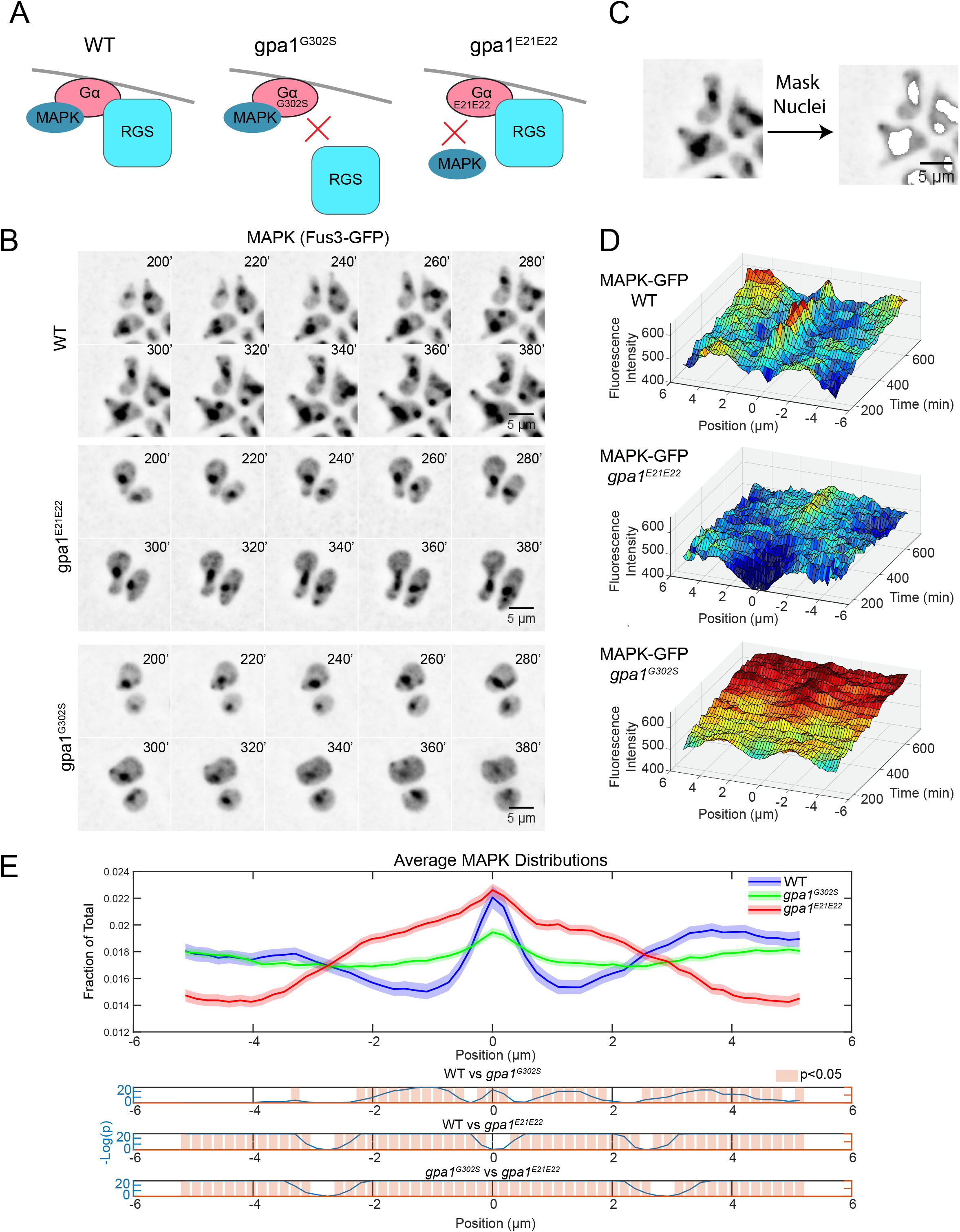
The MAPK Fus3 can serve as an indicator of the localization of active Gα. A) Diagram of the effect of the hyperactive *gpa1^G302S^* mutant and the MAPK-uncoupling *gpa1^E21E22^* mutants. B) Epifluorescence time course images of Fus3-GFP with the indicated Gα mutants. Cells were imaged in a microfluidic device for the indicated time and exposed to a flat 300nM pheromone concentration. C) In order to quantify peripheral MAPK, the nuclear signal was masked for each cell prior to quantitation. D) Average kymographs of MAPK localization in the indicated cell line from (B). E) Quantification of the amount of MAPK on the periphery of the cell spatially normalized to the center of the polar cap as in Figure 2. This analysis is based on the average projection of the stack of images, as that more accurately represents the amount of protein present on the periphery. Shaded areas represent 95% confidence intervals. Data is derived from n = 51 (WT), n = 157 (*gpa1^G302S^)*, and n = 86 (*gpa1^E21E22^)*, with 29 time points per cell.

Having determined that MAPK localization can serve as a proxy for the localization of active Gα and that RGS localization is altered by phosphorylation, we next examined whether RGS phosphorylation affects the distribution of the Gα/MAPK complex. We generated strains expressing fluorescent-protein fusions of a polar cap marker (Bem1-mRuby2) and MAPK (Fus3- EGFP) in the presence of the phospho-mimetic and unphosphorylatable RGS mutants. In both mutants, the presence of MAPK at the periphery was much more pronounced than we observed in WT cells (Figure 5A). This consistent signal is evident in the decreased noise in the kymographs (Figure 5B). Examining the distribution relative to the polar cap over time (Figure 5B) we find that MAPK levels are increasing in both, reminiscent of the hyperactive *gpa1^G302S^*. Despite these similarities, we find differences between the two phospho-mutants over time (Figure 5B), and on average (Figure 5C). When examining the kymographs, we see that the unphosphorylatable pRGS mutant shows a distinct peak at the center with a slight dip peripheral to the peak, in the same way that MAPK in WT cells behaves (Figure 4D, 5C). The unphosphorylatable RGS mutant (*sst2^S539A^*) resulted in a very similar average MAPK distribution to WT (Figure 5C). The p*RGS (*sst2^S539D^*) however, displayed an enrichment of MAPK close to the polar cap (Figure 5C), without the slight depression next to the peak (Figure 5B, C). These data again suggest that unphosporylated RGS is the dominant form in the cell at any given time.

**Figure 5.**
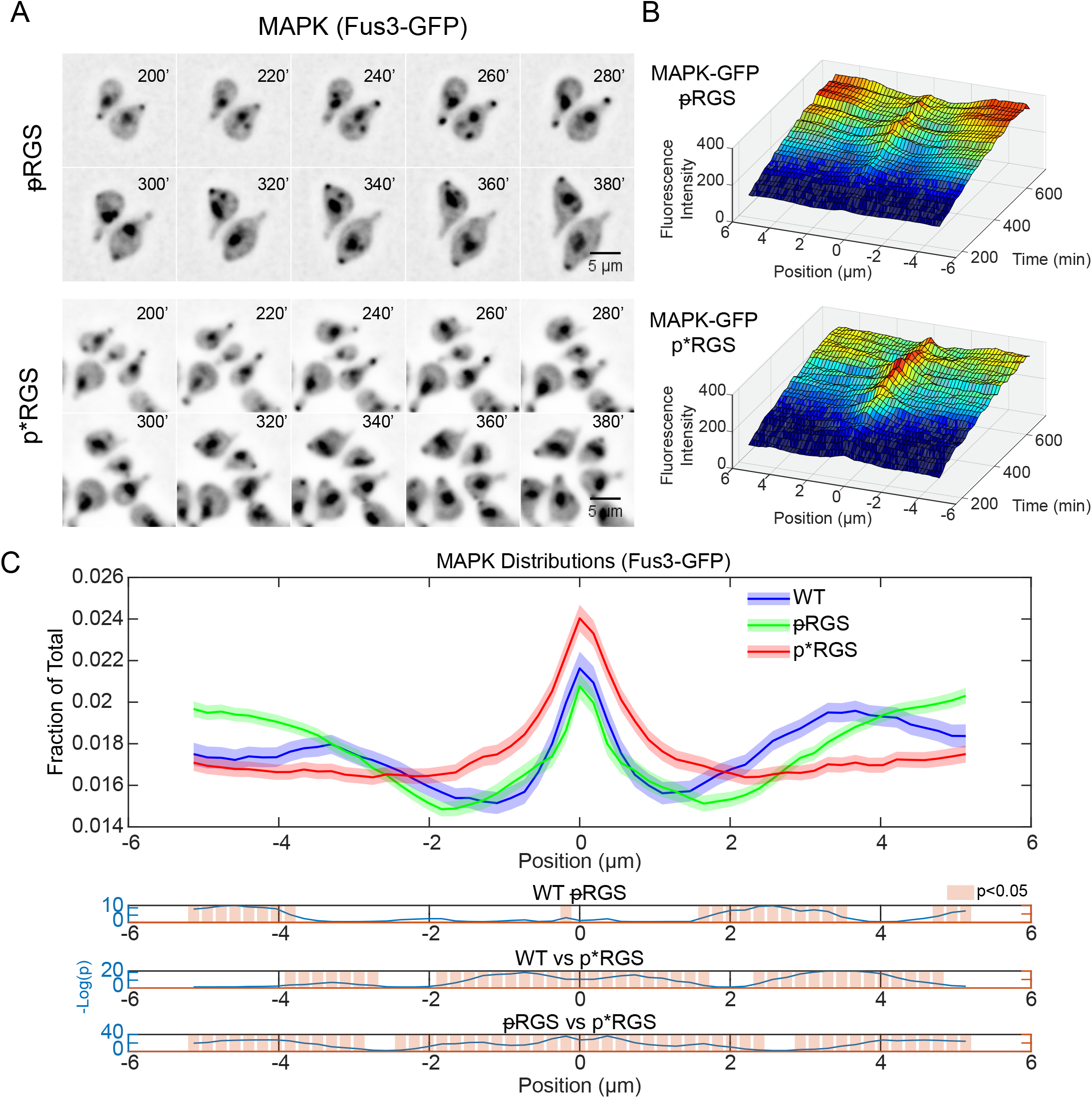
RGS Phosphorylation increase Gα/MAPK complex levels at the center of the polar cap. A) Epifluorescence time course images of Fus3-GFP with the indicated RGS phospho- mutants. Cells were imaged in a microfluidic device for the indicated time and exposed to a flat 300nM pheromone concentration. B) Average kymographs of MAPK localization in the indicated cell line shown in (A). C) Average protein distribution profiles of MAPK (Fus3-GFP) in cells expressing the unphosphorylatable RGS (*sst2^S539A^*) or the phosphomimetic p*RGS (*sst2^S539D^*) aligned to the center of the polar cap (Bem1) as described in Figure 2. The phosphomimetic p*RGS causes a broader distribution of MAPK at the center of the polar cap. Shaded areas represent 95% confidence intervals. Statistical analysis is shown in the graphs below. Data is derived from n = 89(RGS) and n = 73 (p*RGS) cells and 29 time points per cell.

A potential explanation for this increase in average MAPK signal at the polar cap would be a change in the absolute distribution of Gα/MAPK at individual times, or alternatively, the same distribution could have a different relationship with the polar cap. To assess the RGS induced changes in MAPK localization, we examined the distribution of MAPK (Fus3—GFP) spatially normalized to itself instead of Bem1 (Figure 6A). This will show us the shape of MAPK localization at any given time in the cell. We see very little difference in the distribution of MAPK in both phospho-mutants when compared to WT RGS (Figure 6A). Although the changes are statistically significant, they are not large enough to account for the changes in the MAPK localization relative to the polar cap seen in Figure 4. If MAPK has the same shape within the cell under these different conditions, then the offset of this shape relative to the polar cap is changing. To test this, we examined the distribution of the maximum and minimum MAPK intensity relative to maximal polar cap intensity (Figure 6B). The localization of the maxima appears to recapitulate the average distributions in Figure 5C, with the phospho-mimetic p*RGS leading to much more frequent MAPK localization to the polar cap. Perhaps more surprising is the distribution of the minima. In WT cells, the most common place to have a minimum intensity of MAPK is immediately proximal to the center of the polar cap, peaking approximately 1 um away (Figure 6B). In the unphosphorylatable RGS mutant, the minima are again proximal to the center of the polar cap, however they are closer than in WT, peaking at 0.3 to 0.5 µm from the center. In the phosphomimetic p*RGS cells, these minima have largely disappeared, and the minima appear to be much more evenly distributed across the membrane. This suggests an RGS-dependent negative feedback to MAPK proximal to the polar cap that is disrupted by the phosphorylation of the RGS.

**Figure 6.**
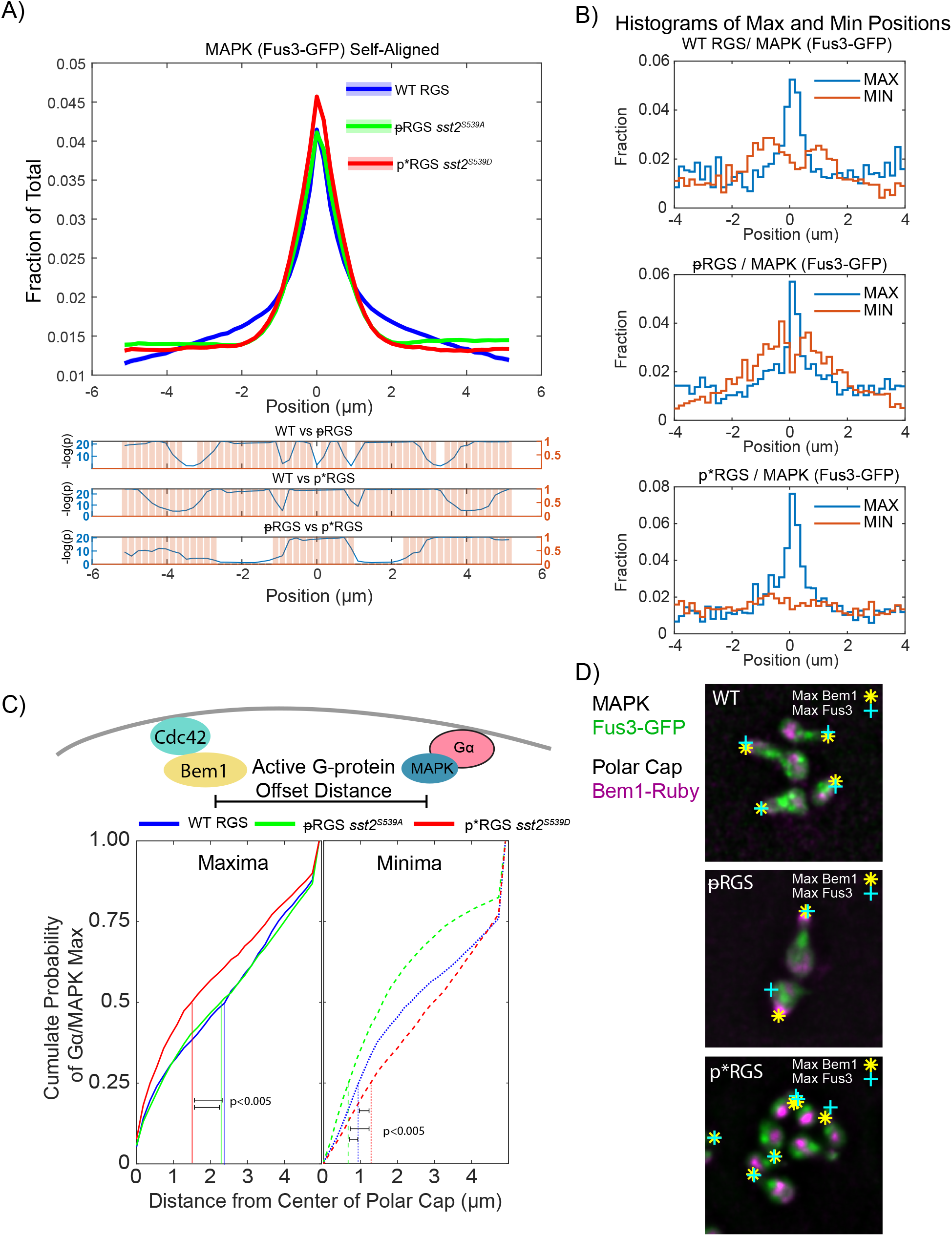
RGS induced changes in Gα/MAPK distribution. A) Distribution of MAPK (Fus3-GFP) from Figure 4, spatially normalized to peak MAPK (Fus3) rather than to the polar cap (Bem1). Shaded areas represent 95% confidence intervals. Statistical analysis is shown in graphs below. B) Histograms of the location of maxima and minima of MAPK spatially registered to the polar cap in the indicated strain. C) We compare the distance between the maxima of the Bem1 and MAPK, effectively the offset between peak Cdc42 activation and peak Gα activation. Graphed is the cumulative sum of MAPK maxima (left) and minima (right) versus distance from the polar cap. Vertical lines show the distance where 50% of maxima have appeared and where 25% of minima have appeared. Statistical significance was evaluated by pairwise Kolomogorov-Smirnov tests. D) Examples of the offset between maximum Bem1 and maximum Fus3 intensity with the indicated RGS mutants.

We then plotted the cumulative distribution of the distance of maxima and minima from the polar cap. This is effectively measuring the offset along the membrane between Bem1 and Fus3, representing active Cdc42 and active Gα. We find that 50% of maxima for WT and RGS fall within ∼ 2.3 µm of the polar cap peak, while 50% of MAPK maxima in the p*RGS fall within ∼1.5 µm of the polar cap peak (p-values calculated using pairwise Kolmogorov-Smirnov tests). We provide some examples of the localizations of the maximum Bem1 and Fus3 in Figure 6D. In examining where there is a large offset between the polar cap and MAPK, it most often appears in those situations where the MAPK intensity at the polar cap is low, and therefore other sites along the periphery may be maximal without a significant accumulation of MAPK. Thus, unphosphorylated RGS drives a greater distance between the polar cap and MAPK, and based on their binding partners, active Gα and active Cdc42.

When examining the minima (Figure 6C) in the unphosphorylatable RGS we have drawn attention to the 25^th^ percentile mark, as approximately 25% of minima in WT occur within 1 µm, corresponding to the WT peak of minima identified in Figure 6B. In the unphosphorylatable RGS mutant, 25% of minima occur within ∼0.6 µm, while in the phospho-mimetic p*RGS, 25% of minima occur within 1.2 µm. The difference in the distributions of the minima are statistically significant between all three strains, and recapitulate our summary of Figure 5B. We conclude that the phosphorylation of the RGS likely disrupts a negative feedback event targeted proximal to the site of polarity.

### RGS phosphorylation peaks early in the pheromone response and diminishes at later time points

Previous characterization of the phosphorylation of the RGS, Sst2, at serine 539 showed Fus3-dependent phosphorylation at one-hour of pheromone treatment (Garrison et al., 1999). By contrast, we examine our cells in microfluidics devices for 12 hours, and we specifically calculate the average distribution of the RGS from 180 minutes onward (approximately the time that the RGS levels reach a new, pheromone-induced steady state, Figure 2C). Our analysis of RGS localization showed that the unphosphorylatable RGS (sst2^S539A^) was similar to WT over the 12-hour time course (Figure 3C). Furthermore, the offset of Gα/MAPK from the polar cap was also similar between the two strains. These observations suggest that unphosphorylated RGS may be the dominant species at later times during the pheromone response. We hypothesized that Sst2 phosphorylation at S539 may be dynamic: peaking earlier in the response and decreasing at later times. To test the dynamics of phosphorylation, we developed a polyclonal antibody to detect Sst2 phosphorylated on serine 539, LHPH**<u>S</u>**PLSEC, where the underlined serine is phosphorylated. Western blotting of Sst2-GFP versus untagged Sst2 shows a GFP-dependent size shift in the detected band, indicating that the antibody is specific for Sst2 (Supplemental Figure 1). Western blotting is done in the presence of excess unphosphorylated peptide to ensure that it is specific for the phospho-epitope.

We next tested whether RGS phosphorylation decreased at later times in the pheromone response. We treated two different strains, *SST2-GFP* and *bar1Δ* with pheromone and took samples every hour for four hours. A complication of examining a potential decrease in pheromone-driven signaling is the desensitizing role of the protease Bar1, which degrades pheromone (Ciejek and Thorner, 1979). By comparing these two strains we can verify that the antibody is detecting Sst2, as the detected band will show a size shift due to the GFP fusion, and we are able to exclude Bar1-mediated degradation of pheromone as driving the decrease in phosphorylation over time. We find that phosphorylation of the RGS peaks between 1 and 2 hours consistent with the literature (Garrison et al., 1999), but phosphorylation levels decrease to lower levels by four hours post pheromone treatment (Figure 7A, B). This decrease is independent of Bar1 activity (Figure 7, Supplemental Figure S1). This decrease is also not due to changes in RGS levels, as RGS is induced in response to pheromone and expression remains high throughout the response (Figure 3D). Decreasing levels of phospho-RGS suggest that the role of the phosphorylated form of the RGS may be more important earlier in the pheromone response.

**Figure 7.**
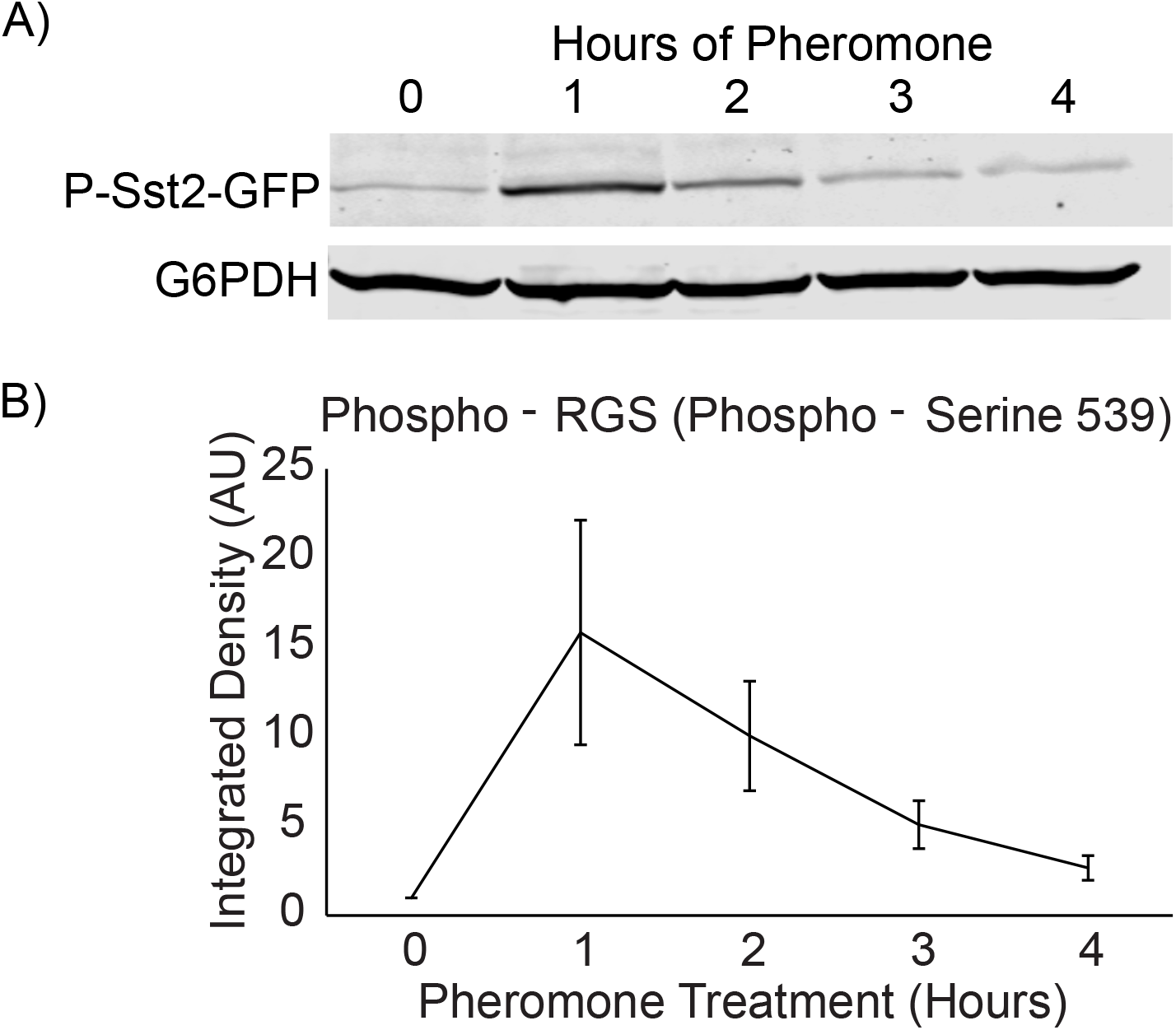
RGS phosphorylation peaks 1hr in the pheromone response. A) Western blotting of phospho-RGS-GFP responding to saturating pheromone over four hours. G6PDH was probed as a loading control. B) Quantification of Western blotting shown in A, normalized to G6PDH levels. Error bars represent standard error of the mean, n=3.

### Phosphorylation of RGS promotes coordination of cytokinesis with the pheromone response

Cells which have already left G1 must complete mitosis and cytokinesis prior to polarizing and forming a mating projection in response to pheromone. We found that some mother-daughter pairs in our unphosphorylatable RGS mutant (*sst2^S539A^*) formed mating projections before they had finished cytokinesis (Figure 8A). The frequency of the event was low and we never observed these defects in wild type cells or cells expressing the phosphomimetic mutant during microfluidics experiments. Previous studies have found interactions between the RGS and two proteins involved in cytokinesis, Bnr1 and Kel1 (Burchett et al., 2002; Yu et al., 2008). Both of these proteins play a role in mitosis, Bnr1 through the regulation of actin polymerization at the mitotic septin ring and Kel1 through promoting the mitotic exit network (MEN) (Buttery et al., 2007; Gao et al., 2010; Hotz and Barral, 2014; Pruyne et al., 2004; Seshan et al., 2002). Additionally, Kel1 serves as a negative regulator of Bnr1, and may impact cytokinesis through that role as well (Gould et al., 2014). Therefore, we hypothesized the cytokinetic defect may be mediated through interactions with these proteins.

**Figure 8.**
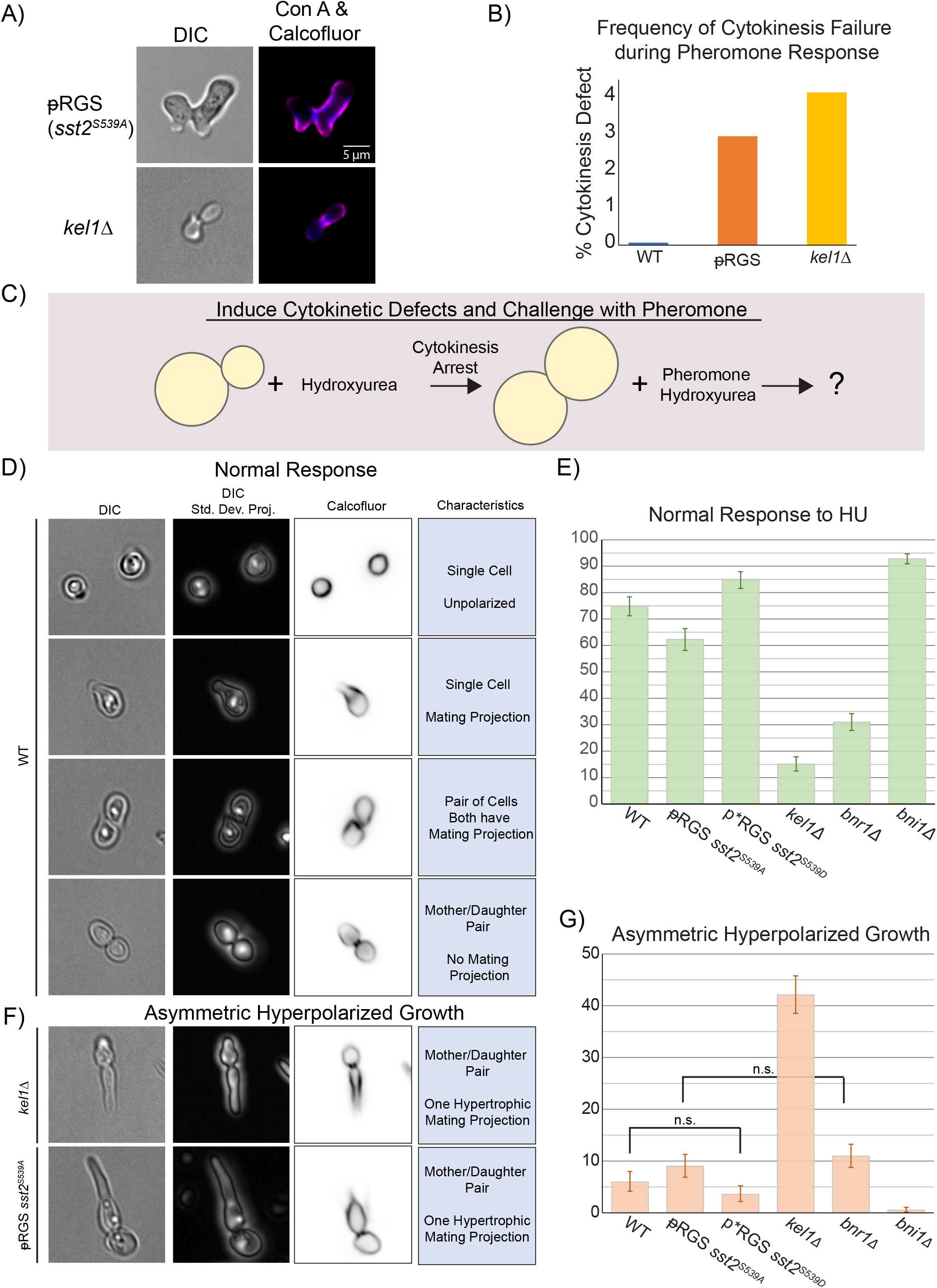
Phosphorylated Sst2 and the Kelch-repeat protein Kel1 promote completion of cytokinesis prior to pheromone induced polarization. A) Images of RGS mutant and *kel1Δ* which have failed to complete cytokinesis prior to pheromone induce polarized growth. Cell walls were stained with Calcofluor White and Concanavalin-A 647 to verify the open bud neck. B) Wild type and mutant RGS and *kel1Δ* strains were exposed to pheromone in culture for 90 minutes, fixed, and then failed cytokinetic events were counted. n = 1412 (WT), 1350 (RGS), and 1396 (*kel1Δ*) from two separate experiments. Total count is shown. C) In order to drive cytokinetic defects, we pretreated cells with 100mM hydroxyurea, followed by treatment with both hydroxyurea and pheromone to investigate the role of the indicated proteins in delay of pheromone induced polarity until completed mitosis. D) Images of normal phenotypes in response to HU + pheromone. Shown are a single focal plane of DIC, a standard deviation projection of a stack of DIC images to better show the state of the bud neck, and cell wall staining with Calcofluor White. We considered a normal response to hydroxyurea and pheromone to be one of the following: 1) Completion of cytokinesis but arrest as a circular cell, in the event that stress signaling is suppressing the pheromone response (a minority of cells). 2) A lone cell responding to pheromone. 3) Completion of cytokinesis (if the cells had resolved their DNA damage), followed by pheromone induced morphogenesis. These cells may be still associated, but show signs of completed cytokinesis 4) Arrest of cytokinesis yielding a mother daughter pair with no polarized growth. E) Plots of the frequency of normal response to hydroxyurea and pheromone in the indicated strains. Error bars represent 95% confidence intervals. For each strain, n > 640 cells across 3 experiments. All differences are statistically significant for p < 0.05, as indicated by non-overlapping 95% confidence intervals. F) Examples of the asymmetric hyperpolarized growth phenotype. G) Plots of the frequency of asymmetric hyperpolarized growth in response to hydroxyurea and pheromone. WT and *sst2^S539A^* have a non-zero difference in means for p < 0.05. Comparisons that are not statistically significant are marked “n.s.”

We examined *bnr1Δ* and *kel1Δ* cells responding to saturating pheromone and found that *kel1Δ* cells also occasionally fail to complete cytokinesis prior to responding to pheromone (Figure 8A, negative data for *bnr1Δ* not shown). These were both rare events and our microfluidic experiments do not contain large numbers of yeast, so we grew wild type, RGS, and *kel1Δ* yeast in culture, treated with saturating pheromone, and counted the frequency of cells with failed cytokinesis based on visual inspection for conjoined yeast responding to pheromone by DIC imaging. We found that both mutants lead to rates of failed cytokinesis of ∼3-4% (Figure 8B, minimum of 1350 cells per strain). From this, we conclude that both Sst2 and Kel1 are both involved in a mechanism that ensures cytokinesis finishes prior to the pheromone response.

To test the role of phosphorylation of the RGS in promoting cytokinesis, we took advantage of hydroxyurea (HU) to damage DNA and cause stalled cytokinesis (Amaral et al., 2016), followed by pheromone exposure, while maintaining the hydroxyurea stress (Figure 8C). HU causes DNA damage to activate the NoCut pathway, an aurora kinase-dependent checkpoint that stalls cytokinesis if chromatin has not cleared the midbody (Amaral et al., 2016; Norden et al., 2006). We reasoned that any process that delays the polarity machinery from being repurposed by pheromone receptor-driven signaling may exacerbate the defects that spontaneously arise in the unphosphorylatable RGS or *kel1Δ* mutants. Under these conditions, stalled cytokinesis would be common, enhancing our ability to differentiate the behaviors of cells with altered Sst2 and Kel1 signaling. We pretreated with 100 mM HU for two hours and then examined cells four hours post pheromone treatment, while maintaining HU. We found that WT cells frequently stalled cytokinesis with two round cells joined at the bud neck. We scored phenotypes as a normal response if the cells showed evidence of completing cytokinesis prior to undergoing polarized growth, or if they arrested as a mother daughter pair with no evidence of polarized growth (Figure 8D, E). We also found cells that had completed cytokinesis and began mating projection formation. In cells with the unphosphorylatable RGS, we found more cells that had both failed cytokinesis and began polarized growth in one or both the mother and daughter cells. A particularly striking phenotype involves one cell remaining round, while the other shows hyper-polarized growth, which we refer to as asymmetric hyperpolarized growth (Figure 8F). In contrast, cells expressing the phospho-mimetic p*RGS mutant (sst2^S539D^) suppress the hyper-polarized growth seen in the unphosphorylatable RGS mutant. Data sets with non-overlapping 95% confidence intervals are statistically significant for p = 0.05 (Figure 8F). Both phospho-mutants have overlapping confidence intervals with the WT, which does not preclude a statistically significant difference. To compare these, we bootstrapped the confidence interval of the difference in means between each phospho- mutant strain and the wild type strain. We then checked whether 0 mean difference fell within the 95% confidence interval. If a difference of 0 falls outside of the 95% confidence interval, we reject the null hypothesis and determine that the difference is statistically significant with a cutoff of p = 0.05. The differences in hyperpolarized growth between WT and the phospho- mimetic p*RGS were not statistically significant, but the increase in hyperpolarized growth in the unphosphorylatable RGS mutant compared to WT was statistically significant.

When we examine *kel1Δ* cells under these conditions, we find that the asymmetric hyperpolarized growth is a dominant phenotype (Figure 8G). Thus, unphosphorylatable RGS partially phenocopies the loss of Kel1 function. This suggests that phosphorylation of RGS promotes a Kel1 dependent mechanism that prevents the mating pathway from commandeering the polarity machinery prior to the completion of cytokinesis. The asymmetric hyperpolarized growth we see in this experiment is consistent with previously published defects seen with perturbation of the Kel1 binding partner, Lte1 (Geymonat et al., 2010) and suggests a defect in Kel1/Lte1 signaling is brought about by unphosphorylated RGS.

Yeast have two formins: Bni1 is associated with the polar cap and is activated by Cdc42 and the Gα/MAPK complex. Bnr1 is associated with mitotic septin structures and has no known role in the pheromone response but is known to be negatively regulated by Kel1 (Gould et al., 2014). Given the central role that formins play in both mitosis and the pheromone response, we hypothesized that the formins may facilitate the coordination of cytokinesis and the beginning of pheromone induced polarized growth. We performed the same experiment as above, inducing cytokinetic defects with hydroxyurea followed by pheromone treatment, and assessed the ability of cells lacking either Bni1 or Bnr1 to prevent pheromone induced polarization prior to the completion of cytokinesis. Deletion of Bni1 largely stopped polarization of cells prior to completion of cytokinesis, and completely abrogated the asymmetric hyper-elongated phenotype (Figure 8E, G). Deletion of Bnr1 resulted in increased polarization prior to the completion of cytokinesis and increased levels of asymmetric hyper-polarized growth (Figure 8E, G). The asymmetric hyper-elongated growth phenotype is clearly dependent upon Bni1 and inhibited by Bnr1. Thus, coordination of pheromone induced polarity with the completion of cytokinesis is promoted by Bnr1 function and antagonized by Bni1.

## Discussion

Here we set out to determine the role of feedback phosphorylation of the RGS, Sst2. We found that the phosphorylation is dynamic through the pheromone response, reaching a maximal level between 1 and 2 hours into the response (Figure 7). We found that phosphorylation of the RGS alters the localization of the RGS relative to the polar cap during the pheromone response and leads to a broadened distribution of the Gα/MAPK complex (Gpa1/Fus3) (Figure 4, 5). Strikingly, cells unable to phosphorylate the RGS sometimes began polarized growth in response to pheromone without waiting for the completion of cytokinesis. By inducing cytokinetic defects with hydroxyurea, we were able to determine that phosphorylation of the RGS enhances the ability of the cell to stall cytokinesis without initiating pheromone induce polarity, thereby correctly integrating both internal stress response and an external morphogenesis response (Figure 8). This coordination appears to use the kelch repeat protein Kel1, and the formin Bnr1.

### The role of RGS phosphorylation

The RGS Sst2 is the primary negative regulator of the pheromone response, a role dominated by its GTPase activating function to turn off the Gα subunit, Gpa1 (Apanovitch et al., 1998; Chasse et al., 2006). The pheromone-induced feedback phosphorylation of the RGS by the MAPK Fus3 is known to have no effect on the pathway sensitivity (Garrison et al., 1999).

These data lead us to pose the question; why is Sst2 phosphorylated? We conclude that the unphosphorylated RGS is important for normal spatial regulation of Gα signaling, but impedes cytokinesis and so the RGS is phosphorylated by MAPK early in the pheromone response to suppress polarity until the completion of cytokinesis (Figure 8).

An obvious question arises from this investigation: Does the RGS plays a role in cytokinesis in the absence of pheromone? There are multiple lines of evidence to suggest that RGS has no role in normal cytokinesis. First, In previous studies on cells lacking the RGS, we have not observed any cytokinetic defects (Kelley et al., 2015). Secondly, baseline Sst2 levels are an order of magnitude higher in haploids than in diploids (de Godoy et al., 2008). If the RGS played an important role in cytokinesis in the absence of pheromone, then haploid and diploid cells would need different mechanisms for regulating cytokinesis, an unlikely scenario.

Our data is consistent with unphosphorylated RGS inhibiting a subset of Kel1 function, as the unphosphorylatable RGS phenocopies the spontaneous failure to complete cytokinesis before mating projection formation that we see in cells lacking Kel1 (Figure 8). We would expect this inhibition of Kel1 to involve stochiometric binding (directly or through an intermediary) and so in the absence of pheromone, where RGS levels are low (Figure 3C and (Dohlman et al., 1996)) there would be very little impact of unphosphorylated RGS on Kel1 activity. We presume that feedback phosphorylation is most important for cells that have just committed to going through the cell cycle when the mating signal starts, as they will have the most accumulation of RGS by the time they reach cytokinesis. In this scenario, a large amount of unphosphorylated RGS present would be able to inhibit Kel1. Phosphorylation of the RGS, however, would prevent its inhibition of Kel1. In the absence of pheromone, the low levels of unphosphorylated RGS would lead to a small impact on Kel1 activity. We conclude that RGS is phosphorylated early in the response to allow normal Kel1 function during the completion of the cell cycle (Figure 8), and dephosporylated at later times to modulate the location of active Gα (Figure 5), likely through modulation of Kel1 function (Figure 8).

### Coordination of the end of cytokinesis with the beginning of receptor mediated morphogenesis

In an unsynchronized population, cells will be evenly distributed through the 90-minute yeast cell cycle. Upon stimulation with pheromone, receptor signaling will immediately begin with subsequent MAPK activation, and downstream phosphorylation of the protein Far1 (Arkowitz, 2009). Far1 serves two purposes in the pheromone response: (1) to inhibit cyclin dependent kinase activity, leading to arrest in G1 (Pope et al., 2014), and (2) to couple the Cdc42 GEF, Cdc24, to free Gβ, thereby promoting polarization to sites of active receptor (Nern and Arkowitz, 1999). The duration of receptor signaling prior to the repurposing of the polar cap will vary depending on where in the cell cycle each cell is when pheromone signaling begins. Thus, some cells may be an hour or more into pheromone signaling prior to completing cytokinesis, while others may be able to immediately start mating projection formation or experience a delay of only a few minutes. A potentially significant difference between these two scenarios is the amount of RGS present in the cell (Figure 3), as SST2 transcription is upregulated by pheromone signaling (Dohlman et al., 1996), and so cells that must delay receptor driven polarity for a long time prior to cytokinesis may be more prone to RGS-induced errors and be more dependent upon MAPK phosphorylation of the RGS.

We have found the delay of receptor driven polarity is promoted by phosphorylation of the RGS and by the kelch-repeat protein, Kel1, which associates with the polar cap, regulates the formin Bnr1, is required for efficient mating, and takes part in the mitotic exit network (MEN) (Gould et al., 2014; Hofken and Schiebel, 2002; Philips and Herskowitz, 1998; Smith and Rose, 2016). Kel1 contributes to the MEN by anchoring the Ras regulator Lte1 to the daughter cell during mitosis (Geymonat et al., 2010; Hotz and Barral, 2014). In addition to promoting mitotic exit, Kel1 and Lte1 have been found to suppress spurious polarization prior to the completion of mitosis, a role that may be separate from their role in MEN (Geymonat et al., 2010). Failure of Lte1 suppression of polarized growth leads to asymmetric hyperpolarized growth very similar to what we see in HU (Figure 8) (Geymonat et al., 2010). Based on our results with HU, it appears that unphosphorylated RGS is disrupting this Lte1 polarity suppression pathway, while phosphorylation of the RGS promotes Lte1 polarity suppression. Geymonat and colleagues described this suppression of polarity in terms of Rsr1/Bud1 driven polarity (Geymonat et al., 2010), which would normally drive the activation of Cdc42 at the insipient bud site for the next round of mitosis (Kozminski et al., 2003). Here we see a block in polarity driven by Gβγ associated Cdc42 GEF (Cdc24) (Butty et al., 1998), which suggests that the ability of Kel1 and Lte1 to inhibit polarity is broadly applicable and may serve further functions in the pheromone response.

### The role of unphosphorylated RGS

At later time points the RGS is dephosphorylated (Figure 7), suggesting a switch in the requirements for RGS as the pheromone response progresses. We found that the localization of the RGS to the center of the polar cap is inhibited by phosphorylation. This phospho-dependent change in localization may seem unintuitive at first glance, as RGS binds to the Receptor at the periphery of the cell, in an interaction required for proper pathway desensitization (Ballon et al., 2006). Disruption of receptor binding with the *sst2^Q304N^* mutant causes supersensitivity of the pheromone pathway (Ballon et al., 2006; Dixit et al., 2014), but phosphorylation of S539 on the RGS has no effect on pathway sensitivity (Garrison et al., 1999). Additionally, receptor binding is inhibited by receptor phosphorylation (Ballon et al., 2006), and so only a subset of the receptor on the cell surface at any given time may be capable of RGS binding. All of this suggests that the change in polar cap localization of the RGS is due to interactions with a protein other than the receptor, and indeed, we do still see a spike in phosphomimetic p*RGS concentration at the center of the polar cap that may represent the receptor associated RGS. Our data suggest that the functional consequence of RGS phosphorylation is altered spatial regulation of the pathway (Figure 9).

**Figure 9.**
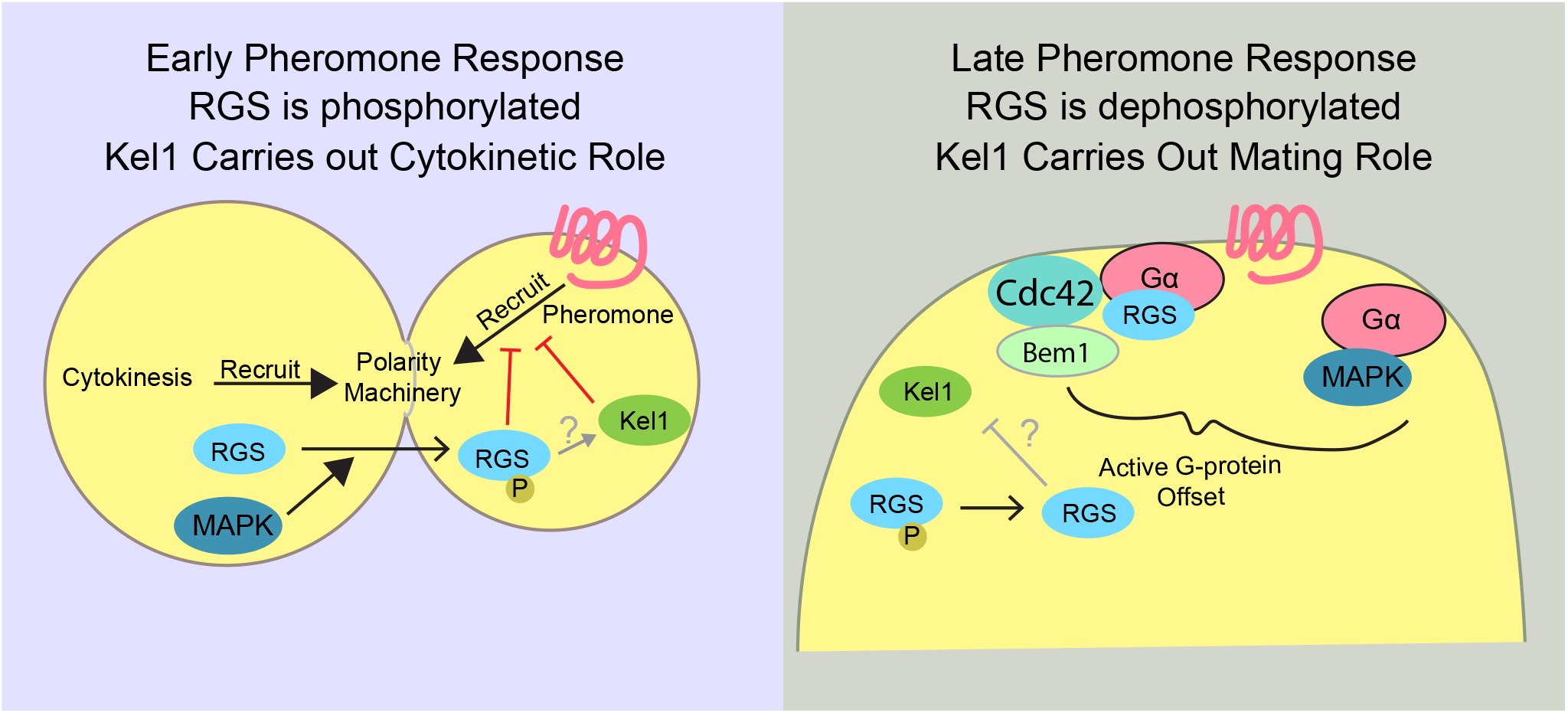
Proposed Role of RGS phosphorylation. In cells that have detected pheromone during mitosis and prior to cytokinesis, phospho-RGS and Kel1 inhibit the recruitment of the polarity machinery by active pheromone receptor. This may be because inactive RGS inhibits Kel1 function. After cytokinesis while cells are generating a mating projection or tracking a gradient of pheromone, unphosphorylated RGS promotes a greater distance between active Cdc42 and active Gα, potentially enhancing polar cap wandering.

Unphosphorylated RGS (e.g. WT at later time points or the RGS mutant) drives a larger distance between active large G-protein (Gpa1) and active small G-protein (Cdc42) than cells expressing the phospho-mimetic p*RGS (Figure 6). The simplest explanation of this observation is that the presence of RGS at the polar cap locally suppresses Gα activation. This is further bolstered by the concentration of minimum MAPK concentration immediately proximal to the center of the polar cap, a phenomenon that is disrupted by the phospho-mimetic mutation that decreases RGS association with the polar cap. This type of negative feedback to the center of active signaling could help drive wandering of the polar cap by promoting large G-protein signaling further from the current site of polarization (Howell et al., 2012). Wandering of the polar cap is important for sensitive gradient tracking (Dyer et al., 2013), and so the small difference in the ability to track the gradient very well that we see in the p*RGS mutant (Figure 1) may be due to its decreased offset between the receptor driven large G-protein and the Cdc42 driven polarity machinery. Additionally, an offset between receptor signaling and the polar cap has recently been proposed to play a role in gradient tracking (Ghose et al., 2021), and while we did not see an effect on gradient tracking here, under more difficult tracking conditions, the observed Gα offset may enhance chemotropic growth.

The phosphorylation of RGS at early time points appears to allow Kel1 to function in delaying early polarization, while unphosphorylatable RGS inhibits that function. This suggests that the role of unphosphorylated RGS at later time points likely interferes with Kel1 function. This may be to regulate the formin, Bnr1, or it may be to regulate Lte1. Regulation of a formin might be the expected function, but Bnr1 appears to have little role in the pheromone pathway. Bni1 would seem a more likely candidate, and while there is no evidence that Kel1 can regulate Bni1, neither is there clear evidence that it cannot (Gould et al., 2014). Our hydroxyurea experiments show that the asymmetric hyper-polarized growth requires Bni1 and is inhibited by Kel1, consistent with a role for Kel1 negatively regulating Bni1, but as mentioned above, this phenotype is likely dependent upon the polarity suppressing effects of Lte1. An intriguing possibility is that unphosphorylated RGS regulates Lte1 function at later times to promote polarity.

## Conclusion

Here we have established a role for the phosphorylation of the RGS, Sst2, in promoting the completion of cytokinesis prior to the pheromone pathway repurposing the mitotic Cdc42 machinery for production of the mating projection, or shmoo. The exacerbation of this defect by the presence of a DNA-damaging agent emphasizes that the cell must integrate the competing signals of a check point instructing the cell to stop mitosis with a GPCR signaling pathway instructing the cell to polarize towards a mating partner. The use of a short-term phosphorylation event to temporarily alter RGS function and thereby allow the mitotic checkpoint to complete seems a likely motif to repeat in other systems.

## Acknowledgements

This research was supported by the National Institute of General Medical Sciences of the National Institutes of Health under Award Number R15GM128026. The content is solely the responsibility of the authors and does not necessarily represent the official views of the National Institutes of Health. The authors declare no conflicts of interest.

## Author Contributions

William Simke contributed to conceptualization, investigation, data curation, formal analysis, investigation, visualization, writing -original draft, and writing -review & editing. Cory Johnson contributed to conceptualization, investigation, methodology, validation, formal analysis and writing -review and editing. Andrew Hart contributed to conceptualization, investigation, formal analysis, and visualization and writing -review and editing. Sari Mayhue contributed to conceptualization, methodology, investigation, and data analysis. Lucas Craig contributed to investigation and formal analysis. Savannah Sojka contributed to investigation. Joshua Kelley contributed to funding acquisition, conceptualization, formal analysis, methodology, investigation, supervision, software, visualization, and writing - original draft.

## Methods

### Yeast Strains

Strains used in this study are shown in Table 2. Strains were constructed in the MATa haploid Saccharomyces cerevisiae strain, BY4741. Proteins were tagged with EGFP or mRuby2 at the native chromosomal locus through oligonucleotide-directed homologous recombination with using the primers listed in Table 3. For tagging Bem1 with Ruby, we created the integrating plasmid pRSII-Bem1-yomRuby2-Kan (table 2). Bem1 nucleic acids 522-1653 were cloned into pRSII405 (Chee and Haase, 2012) followed by link-yomRuby2 from pFA6a-link- yomRuby2 (Lee et al., 2013) using the primers indicated in Table 3. GFP tagging was generated by using pFA6a-link-yoEGFP-spHis5 Kan (Lee et al., 2013) or by amplifying the GFP cassette from the yeast GFP collection (Huh et al., 2003). pFA6a-link-yomRuby2-Kan was a gift from Wendell Lim & Kurt Thorn (Addgene plasmid # 44953 ; http://n2t.net/addgene:44953 ; RRID:Addgene_44953). pRSII405 was a gift from Steven Haase (Addgene plasmid # 35440 ; http://n2t.net/addgene:35440 ; RRID:Addgene_35440).

Sst2 phosphomutants were made by integrating the codon of interest with a PCR amplified CORE cassette(Storici and Resnick, 2006). Deletions were performed by first amplifying the genomic locus from the MATa haploid deletion collection (Dharmacon) with primers listed in Table 3 and transformed using a standard lithium acetate transformation (Burke et al., 2000).

Cells were grown in rich medium (YPD) or synthetic medium (SC) at 30°C unless otherwise indicated. PCR products were transformed into yeast strains using standard lithium acetate transformation procedure. Individual colonies were isolated by growth on standard selective media (SC leu-, SC ura-, SC his-,), selective media with 5-fluoroorotic acid (Zymo Research, Tustin, CA), or YPD selective media (YPD G418+). Transformants were verified using fluorescence microscopy, sequencing, and/or PCR.

### Hydroxyurea experiments

Yeast cultures were grown to an OD600 of 0.4-0.6 at 30°C and then pre-treated with 100 mM hydroxyurea (Alfa Aesar, Tewksbury, MA) for 2 hours at 30°C. After 2 hours of pre-treatment with HU, a saturating concentration of α-factor was added and cultures continued to incubate. Cultures were then fixed at 90 minutes and 240 minutes using an overnight ethanol fixation at -20°C. Following ethanol fixation, yeast were resuspended and washed twice in 50 nM sodium citrate buffer (pH 7.2). Next, the cultures were incubated with 20mg/mL RNase A (Thermo Scientific) for a minimum of 1 hour at 37°C. Following RNase incubation, proteinase K (Thermo Scientific) was added to the cultures at a final concentration of 0.4mg/mL and incubated at 55°C for a minimum of 1 hour then placed at 4°C overnight. For imaging, cells were pelleted, then washed and mounted in 1X PBS (pH 7.4). Cells were then imaged on the IX83 epiflourescent microscope (Olympus).

### Spontaneous cytokinesis defect experiments

Yeast strains were grown in liquid synthetic complete media with 2 % dextrose (SCD) at 30 °C to an OD600 of 0.6 – 0.8. Cells were treated with 30 μM α-factor for 90 minutes. Cells then were fixed in 4 % paraformaldehyde, 2 % glucose, and 30 μM α-factor for 20 minutes. After fixation and 3 washes with 1 X phosphate buffered saline (PBS), the cells were stained with 7 μM Calcofluor White for 20 minutes and 50 μg/mL of Concanavalin A (both obtained from Biotium, Fremont CA) for 30 minutes. The cells were once again washed 3 times with 1x PBS and then imaged. Randomly chosen fields were imaged and then cells were scored for failed cytokinesis.

### Antibody production

The following peptides corresponding to the Sst2 amino acid sequence surround Serine 539 were synthesized by Genscript (Piscataway NJ), the phospho-Sst2 S539 peptide LHPH**<u>S</u>**PLSEC, where the **<u>S</u>** was phosphorylated, and the unphosphorylated peptide LHPHSPLSEC. The phospho-peptide was injected into rabbits by Cocalico Biologicals (Reamstown PA) according to their standard protocol. Antibody was affinity purified on phospho-peptide covalently bound to a SulfoLink column according to the manufacturer’s instructions (Thermo Scientific).

### Western blotting

Phosphorylation state of Sst2 was assessed by Western blotting. Yeast cultures were grown overnight in 30°C. Cells were lysed with TCA buffer and protein concentrations were determined using DC protein assay kit (BioRad). Protein separation was performed with a 7.5% SDS-PAGE and transferred to nitrocellulose at 100 V for 90 mins.

Primary antibody (1:1,000) and non-phosphopepetide (1:10,000) were incubated in 1% PBST blocking solution overnight followed by secondary antibody incubation (1:10,000) in 1% PBST blocking solution for 1 hr. Band intensity was detected via Odyssey CLx imaging system (LI-COR) and then quantified using ImageJ.

### Imaging on Agarose Pad

Yeast were imaged on an Olympus IX83 with a 60X-TIRF 1.49 NA objective, a Photometrics Prime95b camera, X-Cite LED 120 Boost fluorescence light source (Excelitas), and filters for DAPI and GFP (Semrock). Cells were grown to mid-log phase (OD600 = 0.1 to 0.8) at 30°C in Synthetic Complete Media with 2% dextrose (SCD) and then imaged on pads made of 2% agarose in SCD, as the use of agarose leads to lower autofluorescence than standard agar pads. Imaging was performed with an objective heater (Bioptechs) set to 30°C. Cells were pelleted and then resuspended in SC with 3uM α-factor and placed on an agarose pad as above.

### Microfluidics Experiments

Microfluidic devices were made by using a Silicone polymer poured onto a microfluidics device mold (Suzuki et al., 2021) fabricated by UMaine FIRST. SYLGARD 184 Silicone Polymer was mixed at a ratio of 10:1, part A to part B, using a glass stirring rod to mix (Dow). Mixed polymer was poured onto the device mold and placed in a vacuum chamber for 1hr. After all air bubbles were removed, the mixture was placed in an oven at 80°C for 1hr.

After cooling to room temperature, devices were cut out using a razor and ports were punctured using an 18g Leuer stub. Prepped devices and coverslips were cleaned by spraying with methanol, ethanol, then water, and dried using an air hose. Devices and coverslips were exposed to oxygen plasma for 45 seconds in a Harrick Plasma PDC32G Cleaner followed by fusion of the device to the cover slip.

Cultures were grown in SC to an OD600 between 0.1–0.8 at 30°C. Live-cell microfluidics experiments were performed using an IX83 (Olympus, Waltham MA) microscope with a Prime 95B CMOS Camera (Photometrics) controlled by Cell Sens 1.17 (Olympus). Fluorescence and Differential Interference Contrast (DIC) images were acquired using an Olympus-APON-60X-TIRF objective. Z-stacks of GFP and RFP images were acquired using an X-Cite 120 LEDBoost (Excelitas). Cells were imaged in a microfluidic device based on the Dial-a-wave design that allows for the rapid switching of media while holding the yeast in place (Bennett 2008, Dixit 2014)(Suzuki et al., 2021). Pheromone addition was verified using AlexaFluor 647 dye (Life Technologies) imaged with a single plane image. Cells were imaged at 20 min intervals for 12 hours for 300nM experiments and 5 min intervals for 0-150nM experiments. Confocal microscopy was conducted on a Leica DMi8 (Leica) imaging platform equipped with an automated stage, SP8X white light laser (capped at 70% of total power), an argon laser (Leica Microsystems Buffalo Grove, IL). All imaging was conducted using HyD hybrid detectors.

Imaging settings were determined based on experimental needs and were replicated for repeat experiments.

### Image Analysis

Images were deconvolved using Huygens Software (Scientific Volume Imaging, Hilversum, Netherlands) Classic Maximum Likelihood Estimation (CMLE) Deconvolution Algorithm. Masks of cells were made using ImageJ (Schindelin et al., 2012) and data analysis was performed using MATLAB (MathWorks, Natick, MA). To quantify the fraction of protein localization over time, MATLAB was utilized as described in Figure 2 and previously (Kelley et al., 2015; Shellhammer et al., 2019). The fluorescent intensity of each fluorescent protein was extracted over time using a line width of 5 pixels. Peak Bem1 was used as a reference to normalize the spatial distribution of proteins of interest in relation to the polar cap. This was done by setting peak Bem1 as the midpoint and shifting the protein of interest in the same manner. For profiles reporting fraction of protein at each position, fluorescence was normalized by subtracting the minimum value from each line-scan, followed by normalization of the subtracted data to sum to one. The normalized fluorescence intensity was plotted at each point along the cell periphery with shaded regions showing 95% confidence intervals derived by bootstrapping with 10000 resamplings. Statistical analysis was performed between profiles using a sliding one-way ANOVA and Tukeys honestly significant difference (HSD) test followed by false discover rate adjustment with the MATLAB *mafdr()* function with p values <0.05 denoted as significant. Where indicated, a pairwise Kolmogorov-Smirnov test was performed using the MATLAB *kstest2()* function. When excluding nuclear fluorescence from Fus3-GFP images, we modified the “granulinator” script (Hunn et al., 2021) to select nuclei for removal. We used cell masks to calculate fluorescence histograms for each cell and adjusted the size (minimum size of 25 pixels) and threshold (1 standard deviation above mean) cutoffs to detect nuclei but not polar caps.

---

**Supplemental Figure S1.**
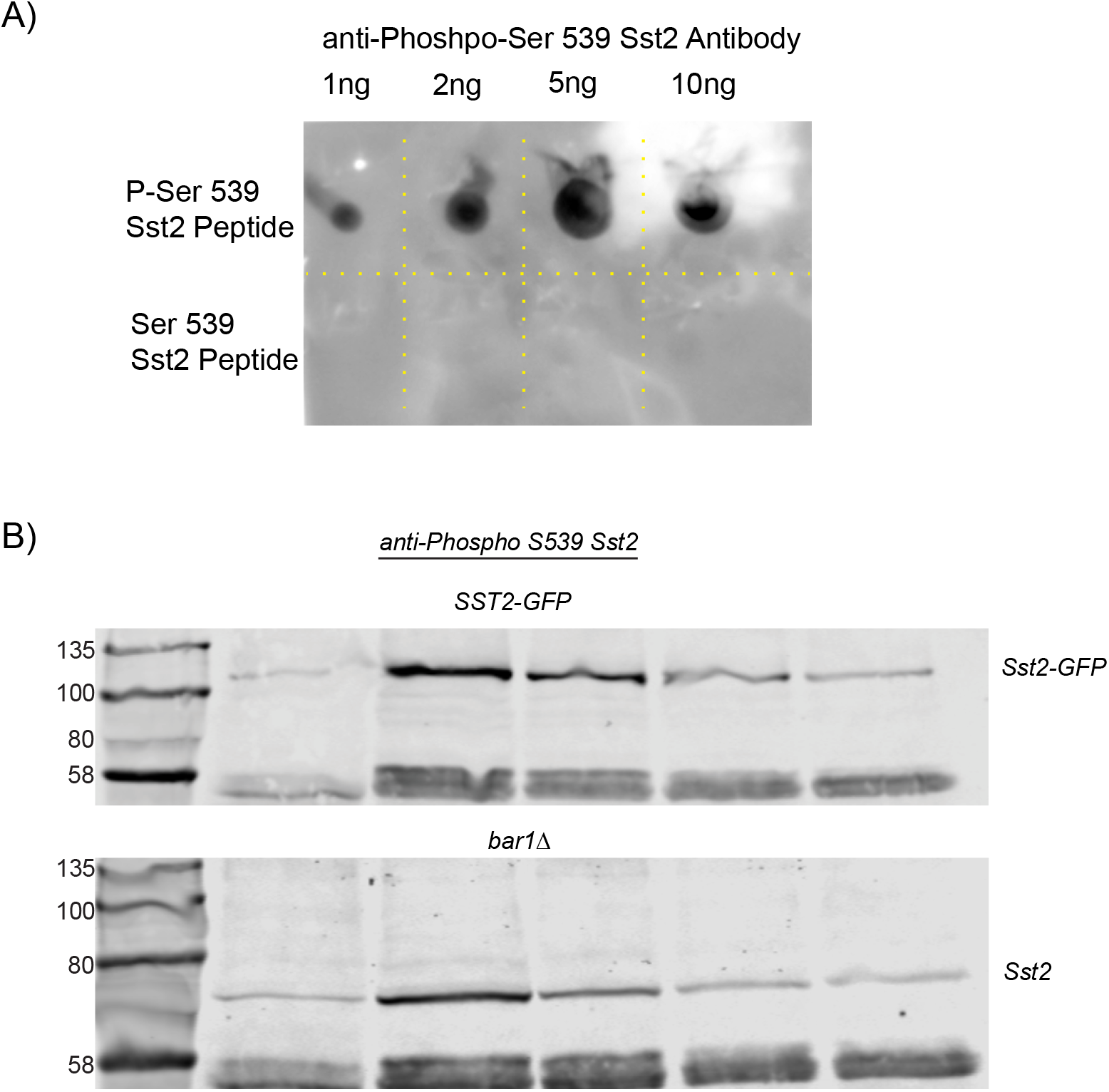
Characterization of the phospho-Sst2 antibody. A) Dot blot with the indicated quantities of peptide. The antibody is specific for phospho-peptide under the conditions used. All blotting is carried out with excess unphosporylated peptide to stop nonspecific binding. B) Our anti-phospo-Sst2 antibody detects a higher molecular weight band when detecting Sst2-GFP than when detecting native Sst2. Additionally, the time dependent decrease in phospho-Sst2 is not dependent upon Bar1-mediated degradation of pheromone.

**Table S1.**

| STRAIN | PARENT | DESCRIPTION |
| --- | --- | --- |
| BY4741 |  | *Mata leu2Δ met15Δ his3Δ ura3Δ* |
| SST2-GFP | BY4741 | *SST2-GFP::HIS3* |
| SST2-GFP BEM1-RUBY2 | BY4741 | *SST2-GFP::HIS3 BEM1-RUBY2::KanMX4* |
| SST2<sup>S539A</sup> | BY4741 | *sst2<sup>S539A</sup>* |
| SST2<sup>S539D</sup> | BY4741 | *sst2<sup>S539D</sup>* |
| SST2<sup>S539A</sup>-GFP BEM1-RUBY2 | BY4741 | *sst2<sup>S539A</sup>-GFP::URA3 BEM1-RUBY2::KanMX4* |
| SST2<sup>S539D</sup>-GFP | BY4741 | *sst2<sup>S539D</sup>-GFP::HIS3* |
| SST2<sup>S539D</sup>-GFP BEM1-RUBY2 | BY4741 | *sst2<sup>S539D</sup>-GFP::URA3 BEM1-RUBY2::LEU2* |
| KEL1Δ | BY4741 | *kel1Δ::KanMX4* |
| GPA1<sup>G302S</sup> | BY4741 | *gpa1<sup>G302S</sup>::URA3* |
| GPA1<sup>G302S</sup> SST2-GFP BEM1-RUBY2 | BY4741 | *gpa1<sup>G302S</sup>::URA3 SST2-GFP::HIS3 Bem1-RUBY2::LEU2* |
| GPA1<sup>EE</sup> SST2-GFP BEM1-RUBY2 | BY4741 | *gpa1<sup>EE</sup>::URA3 SST2-GFP::HIS3 Bem1-RUBY2::LEU2* |
| GPA1<sup>EE</sup> FUS3-GFP BEM1-RUBY2 | BY4741 | *gpa1<sup>EE</sup>::URA3 FUS3-GFP::HIS3 Bem1-RUBY2::LEU2* |
| BNI1Δ | BY4741 | *bni1Δ::KanMX4* |
| BNR1Δ | BY4741 | *bnr1Δ::KanMX4* |
| FUS3-GFP BEM1-RUBY2 | BY4741 | *FUS3-GFP::HIS3 BEM1-RUBY2::LEU2* |
| SST2<sup>S539A</sup> FUS3-GFP BEM1-RUBY2 | BY4741 | *sst2<sup>S539A</sup> FUS3-GFP::HIS3 BEM1-RUBY2::LEU2* |
| SST2<sup>S539D</sup> FUS3-GFP BEM1-RUBY2 | BY4741 | *sst2<sup>S539D</sup> FUS3-GFP::HIS3 BEM1-RUBY2::LEU2* |

**Table S2.**

| PLASMID | VECTOR | DESCRIPTION |
| --- | --- | --- |
| PRSII405-BEM1-RUBY2 | pRSII405 | Integrating BEM1-RUBY::LEU2 Vector |
| PRSII406-GPA1<sup>G302S</sup> | pRSII406 | Integrating *gpa1<sup>G302S</sup>::URA3* Vector |
| PRS406-SST2-GFP | pRSII406 | Integrating SST2-GFP::URA3 Vector |
| PRSII406-SST2<sup>S539A</sup>-GFP | pRSII406 | Integrating *sst2<sup>S539A</sup>-GFP::URA3* Vector |

**Table S3.**
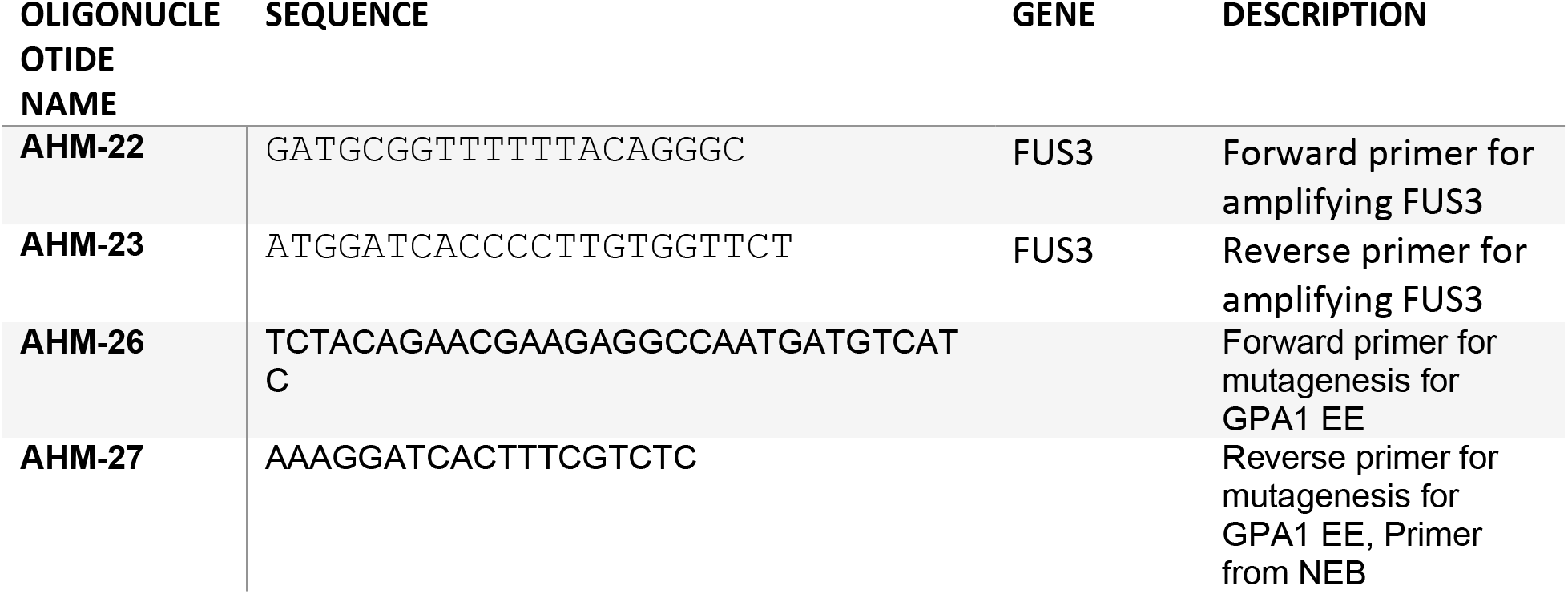
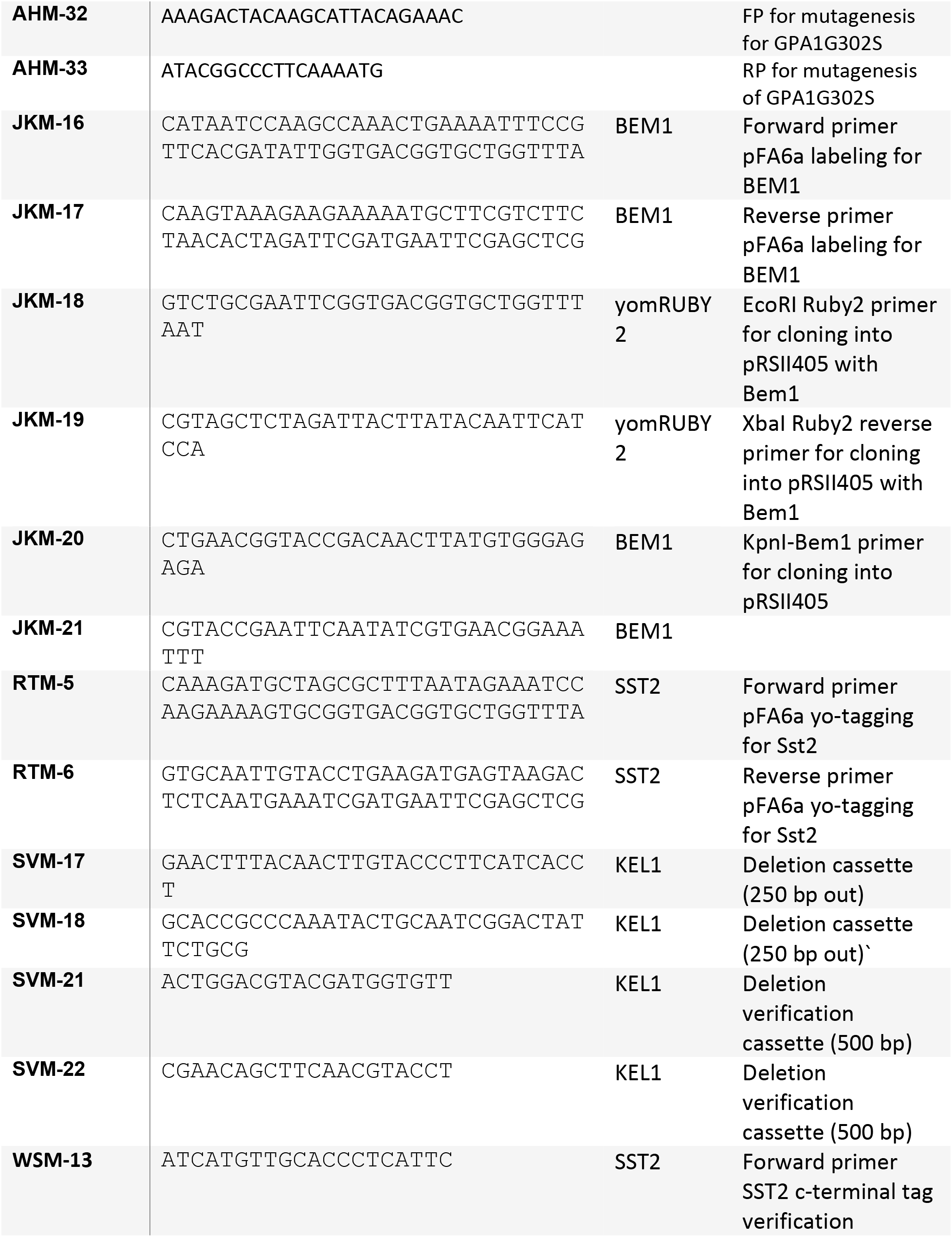
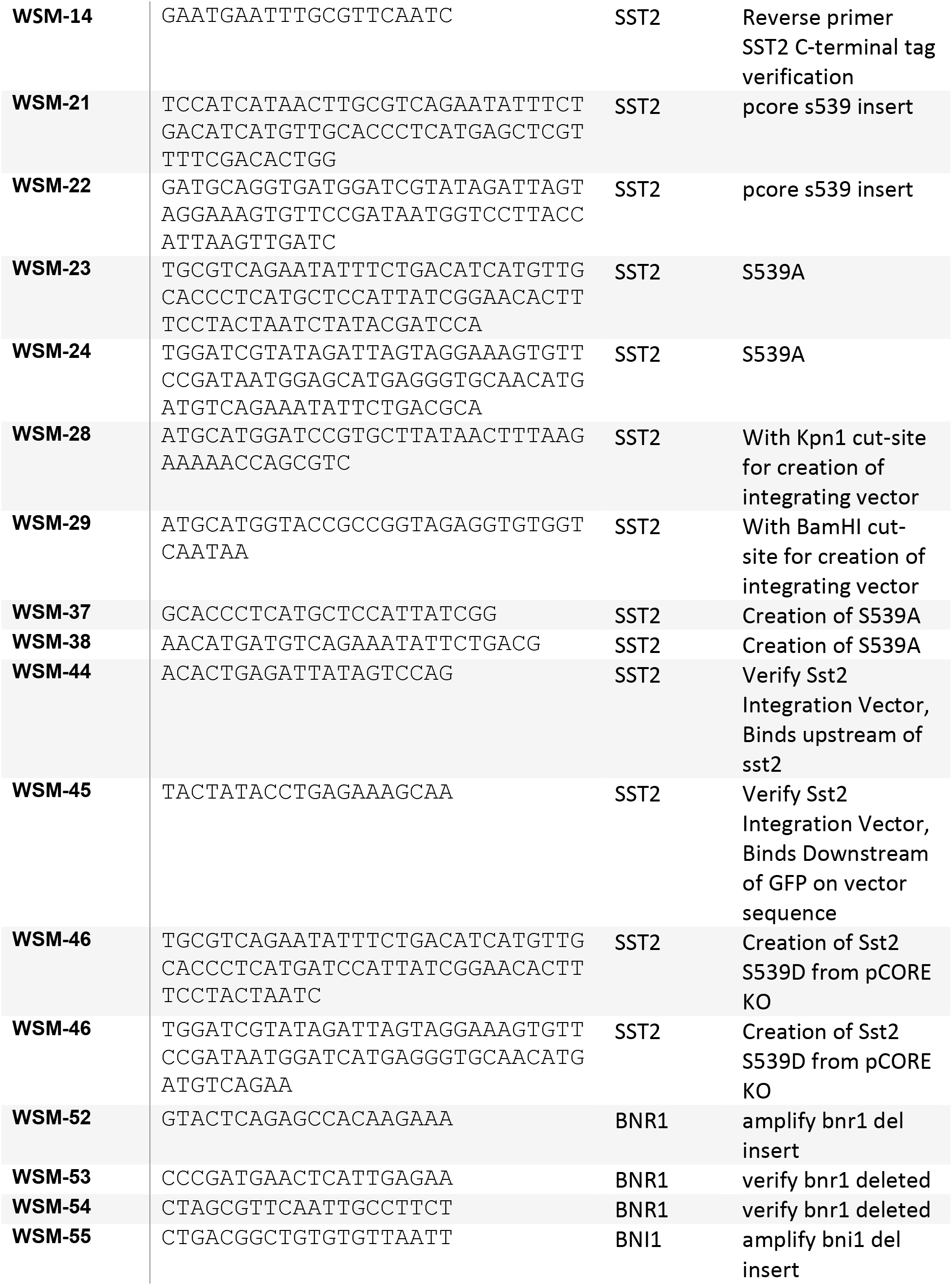
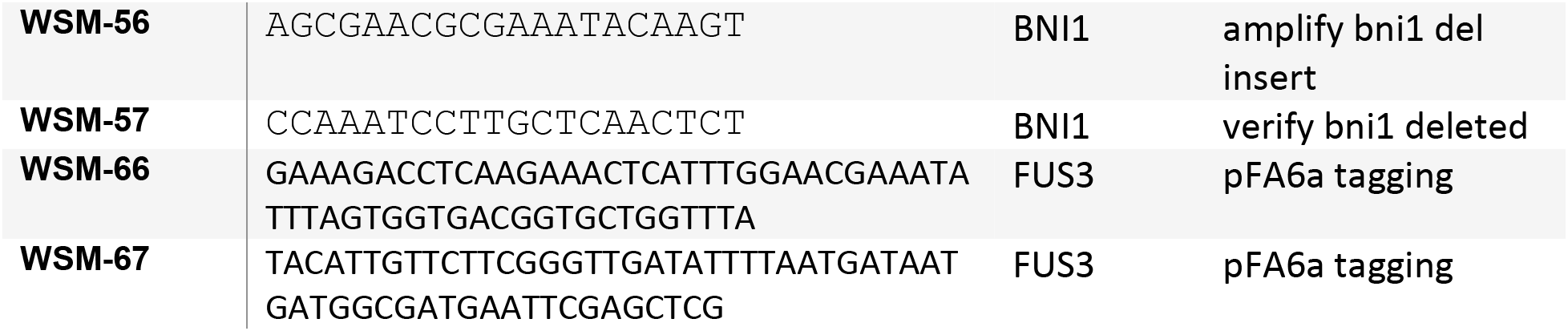

## References

Alvaro, C.G., and J. Thorner. 2016. Heterotrimeric G Protein-coupled Receptor Signaling in Yeast Mating Pheromone Response. The Journal of biological chemistry. 291:7788–7795.

Amaral, N., A. Vendrell, C. Funaya, F.Z. Idrissi, M. Maier, A. Kumar, G. Neurohr, N. Colomina, J. Torres-Rosell, M.I. Geli, and M. Mendoza. 2016. The Aurora-B-dependent NoCut checkpoint prevents damage of anaphase bridges after DNA replication stress. Nat Cell Biol. 18:516–526.

Apanovitch, D.M., K.C. Slep, P.B. Sigler, and H.G. Dohlman. 1998. Sst2 is a GTPase-activating protein for Gpa1: purification and characterization of a cognate RGS-Galpha protein pair in yeast. Biochemistry. 37:4815–4822.

Arkowitz, R.A. 2009. Chemical gradients and chemotropism in yeast. Cold Spring Harbor perspectives in biology. 1:a001958.

Ballon, D.R., P.L. Flanary, D.P. Gladue, J.B. Konopka, H.G. Dohlman, and J. Thorner. 2006. DEP-domain-mediated regulation of GPCR signaling responses. Cell. 126:1079–1093.

Bar-Shavit, R., M. Maoz, A. Kancharla, J.K. Nag, D. Agranovich, S. Grisaru-Granovsky, and B. Uziely. 2016. G Protein-Coupled Receptors in Cancer. Int J Mol Sci. 17.

Bi, E., and H.O. Park. 2012. Cell polarization and cytokinesis in budding yeast. Genetics. 191:347–387.

Breitsprecher, D., and B.L. Goode. 2013. Formins at a glance. J Cell Sci. 126:1–7.

Burchett, S., P. Flanary, C. Aston, L. Jiang, K. Young, P. Uetz, S. Fields, and H. Dohlman. 2002. Regulation of stress response signaling by the N-terminal dishevelled/EGL- 10/pleckstrin domain of Sst2, a regulator of G protein signaling in Saccharomyces cerevisiae. The Journal of biological chemistry. 277:22156–22167.

Burke, D., D. Dawson, and T. Stearns. 2000. Methods in Yeast Genetics: A Cold Spring Harbor Laboratory Course Manual.

Buttery, S.M., S. Yoshida, and D. Pellman. 2007. Yeast formins Bni1 and Bnr1 utilize different modes of cortical interaction during the assembly of actin cables. Mol Biol Cell. 18:1826–1838.

Butty, A.C., P.M. Pryciak, L.S. Huang, I. Herskowitz, and M. Peter. 1998. The role of Far1p in linking the heterotrimeric G protein to polarity establishment proteins during yeast mating. Science. 282:1511–1516.

Chasse, S., P. Flanary, S. Parnell, N. Hao, J. Cha, D. Siderovski, and H. Dohlman. 2006. Genome-scale analysis reveals Sst2 as the principal regulator of mating pheromone signaling in the yeast Saccharomyces cerevisiae. Eukaryotic cell. 5:330–346.

Chee, M.K., and S.B. Haase. 2012. New and Redesigned pRS Plasmid Shuttle Vectors for Genetic Manipulation of Saccharomycescerevisiae. G3 (Bethesda). 2:515–526.

Chiou, J.G., M.K. Balasubramanian, and D.J. Lew. 2017. Cell Polarity in Yeast. Annu Rev Cell Dev Biol. 33:77–101.

Ciejek, E., and J. Thorner. 1979. Recovery of S. cerevisiae a cells from G1 arrest by alpha factor pheromone requires endopeptidase action. Cell. 18:623–635.

de Godoy, L.M., J.V. Olsen, J. Cox, M.L. Nielsen, N.C. Hubner, F. Frohlich, T.C. Walther, and M. Mann. 2008. Comprehensive mass-spectrometry-based proteome quantification of haploid versus diploid yeast. Nature. 455:1251–1254.

DiBello, P.R., T.R. Garrison, D.M. Apanovitch, G. Hoffman, D.J. Shuey, K. Mason, M.I. Cockett, and H.G. Dohlman. 1998. Selective uncoupling of RGS action by a single point mutation in the G protein alpha-subunit. The Journal of biological chemistry. 273:5780–5784.

Dixit, G., J.B. Kelley, J.R. Houser, T.C. Elston, and H.G. Dohlman. 2014. Cellular noise suppression by the regulator of G protein signaling Sst2. Molecular cell. 55:85–96.

Dohlman, H., D. Apaniesk, Y. Chen, J. Song, and D. Nusskern. 1995. Inhibition of G-protein signaling by dominant gain-of-function mutations in Sst2p, a pheromone desensitization factor in Saccharomyces cerevisiae. Molecular and cellular biology. 15:3635–3643.

Dohlman, H.G., J. Song, D. Ma, W.E. Courchesne, and J. Thorner. 1996. Sst2, a negative regulator of pheromone signaling in the yeast Saccharomyces cerevisiae: expression, localization, and genetic interaction and physical association with Gpa1 (the G-protein alpha subunit). Molecular and cellular biology. 16:5194–5209.

Dyer, J.M., N.S. Savage, M. Jin, T.R. Zyla, T.C. Elston, and D.J. Lew. 2013. Tracking shallow chemical gradients by actin-driven wandering of the polarization site. Current biology : CB. 23:32–41.

Elion, E.A., B. Satterberg, and J.E. Kranz. 1993. FUS3 phosphorylates multiple components of the mating signal transduction cascade: evidence for STE12 and FAR1. Mol Biol Cell. 4:495–510.

Errede, B., L. Vered, E. Ford, M.I. Pena, and T.C. Elston. 2015. Pheromone-induced morphogenesis and gradient tracking are dependent on the MAPK Fus3 binding to Galpha. Mol Biol Cell. 26:3343–3358.

Evangelista, M., K. Blundell, M.S. Longtine, C.J. Chow, N. Adames, J.R. Pringle, M. Peter, and C. Boone. 1997. Bni1p, a yeast formin linking cdc42p and the actin cytoskeleton during polarized morphogenesis. Science. 276:118–122.

Gao, L., W. Liu, and A. Bretscher. 2010. The yeast formin Bnr1p has two localization regions that show spatially and temporally distinct association with septin structures. Mol Biol Cell. 21:1253–1262.

Garrison, T., Y. Zhang, M. Pausch, D. Apanovitch, R. Aebersold, and H. Dohlman. 1999. Feedback phosphorylation of an RGS protein by MAP kinase in yeast. The Journal of biological chemistry. 274:36387–36391.

Geymonat, M., A. Spanos, S. Jensen, and S.G. Sedgwick. 2010. Phosphorylation of Lte1 by Cdk prevents polarized growth during mitotic arrest in S. cerevisiae. The Journal of cell biology. 191:1097–1112.

Ghose, D., K. Jacobs, S. Ramirez, T. Elston, and D. Lew. 2021. Chemotactic movement of a polarity site enables yeast cells to find their mates. Proc Natl Acad Sci U S A. 118.

Gould, C.J., M. Chesarone-Cataldo, S.L. Alioto, B. Salin, I. Sagot, and B.L. Goode. 2014. Saccharomyces cerevisiae Kelch proteins and Bud14 protein form a stable 520-kDa formin regulatory complex that controls actin cable assembly and cell morphogenesis. The Journal of biological chemistry. 289:18290–18301.

Hao, N., S. Nayak, M. Behar, R.H. Shanks, M.J. Nagiec, B. Errede, J. Hasty, T.C. Elston, and H.G. Dohlman. 2008. Regulation of cell signaling dynamics by the protein kinase-scaffold Ste5. Molecular cell. 30:649–656.

Hauser, A.S., M.M. Attwood, M. Rask-Andersen, H.B. Schioth, and D.E. Gloriam. 2017. Trends in GPCR drug discovery: new agents, targets and indications. Nat Rev Drug Discov. 16:829–842.

Hofken, T., and E. Schiebel. 2002. A role for cell polarity proteins in mitotic exit. The EMBO journal. 21:4851–4862.

Hotz, M., and Y. Barral. 2014. The Mitotic Exit Network: new turns on old pathways. Trends Cell Biol. 24:145–152.

Howell, A., M. Jin, C.-F. Wu, T. Zyla, T. Elston, and D. Lew. 2012. Negative feedback enhances robustness in the yeast polarity establishment circuit. Cell. 149:322–333.

Huh, W.K., J.V. Falvo, L.C. Gerke, A.S. Carroll, R.W. Howson, J.S. Weissman, and E.K. O’Shea. 2003. Global analysis of protein localization in budding yeast. Nature. 425:686–691.

Hunn, J.C., K.M. Hutchinson, J.B. Kelley, and D. Reines. 2021. An Algorithm to Quantify Inducible Protein Condensates In Eukaryotic Cells. bioRxiv:2021.2008.2026.457826.

Imamura, H., K. Tanaka, T. Hihara, M. Umikawa, T. Kamei, K. Takahashi, T. Sasaki, and Y. Takai. 1997. Bni1p and Bnr1p: downstream targets of the Rho family small G-proteins which interact with profilin and regulate actin cytoskeleton in Saccharomyces cerevisiae. The EMBO journal. 16:2745–2755.

Kelley, J.B., G. Dixit, J.B. Sheetz, S.P. Venkatapurapu, T.C. Elston, and H.G. Dohlman. 2015. RGS proteins and septins cooperate to promote chemotropism by regulating polar cap mobility. Current biology : CB. 25:275–285.

Kozminski, K., L. Beven, E. Angerman, A. Tong, C. Boone, and H. Park. 2003. Interaction between a Ras and a Rho GTPase couples selection of a growth site to the development of cell polarity in yeast. Molecular biology of the cell. 14:4958–4970.

Lappano, R., and M. Maggiolini. 2012. GPCRs and cancer. Acta Pharmacol Sin. 33:351–362.

Lee, S., W.A. Lim, and K.S. Thorn. 2013. Improved Blue, Green, and Red Fluorescent Protein Tagging Vectors for S. cerevisiae. PLoS ONE. 8:e67902.

Matheos, D., M. Metodiev, E. Muller, D. Stone, and M.D. Rose. 2004. Pheromone-induced polarization is dependent on the Fus3p MAPK acting through the formin Bni1p. The Journal of cell biology. 165:99–109.

Metodiev, M.V., D. Matheos, M.D. Rose, and D.E. Stone. 2002. Regulation of MAPK function by direct interaction with the mating-specific Galpha in yeast. Science. 296:1483–1486.

Nern, A., and R.A. Arkowitz. 1999. A Cdc24p-Far1p-Gbetagamma protein complex required for yeast orientation during mating. The Journal of cell biology. 144:1187–1202.

Norden, C., M. Mendoza, J. Dobbelaere, C.V. Kotwaliwale, S. Biggins, and Y. Barral. 2006. The NoCut pathway links completion of cytokinesis to spindle midzone function to prevent chromosome breakage. Cell. 125:85–98.

Park, H.O., and E. Bi. 2007. Central roles of small GTPases in the development of cell polarity in yeast and beyond. Microbiology and molecular biology reviews : MMBR. 71:48–96.

Peter, M., A. Gartner, J. Horecka, G. Ammerer, and I. Herskowitz. 1993. FAR1 links the signal transduction pathway to the cell cycle machinery in yeast. Cell. 73:747–760.

Philips, J., and I. Herskowitz. 1998. Identification of Kel1p, a kelch domain-containing protein involved in cell fusion and morphology in Saccharomyces cerevisiae. The Journal of cell biology. 143:375–389.

Pope, P.A., S. Bhaduri, and P.M. Pryciak. 2014. Regulation of cyclin-substrate docking by a G1 arrest signaling pathway and the Cdk inhibitor Far1. Current biology : CB. 24:1390–1396.

Pruyne, D., L. Gao, E. Bi, and A. Bretscher. 2004. Stable and dynamic axes of polarity use distinct formin isoforms in budding yeast. Mol Biol Cell. 15:4971–4989.

Schindelin, J., I. Arganda-Carreras, E. Frise, V. Kaynig, M. Longair, T. Pietzsch, S. Preibisch, C. Rueden, S. Saalfeld, B. Schmid, J.Y. Tinevez, D.J. White, V. Hartenstein, K. Eliceiri, P. Tomancak, and A. Cardona. 2012. Fiji: an open-source platform for biological-image analysis. Nature methods. 9:676–682.

Segall, J.E. 1993. Polarization of yeast cells in spatial gradients of alpha mating factor. Proc Natl Acad Sci U S A. 90:8332–8336.

Seshan, A., A.J. Bardin, and A. Amon. 2002. Control of Lte1 localization by cell polarity determinants and Cdc14. Current biology : CB. 12:2098–2110.

Shellhammer, J.P., A.E. Pomeroy, Y. Li, L. Dujmusic, T.C. Elston, N. Hao, and H.G. Dohlman. 2019. Quantitative analysis of the yeast pheromone pathway. Yeast. 36:495–518.

Smith, J.A., and M.D. Rose. 2016. Kel1p Mediates Yeast Cell Fusion Through a Fus2p- and Cdc42p-Dependent Mechanism. Genetics. 202:1421–1435.

Storici, F., and M.A. Resnick. 2006. The delitto perfetto approach to in vivo site-directed mutagenesis and chromosome rearrangements with synthetic oligonucleotides in yeast. Methods Enzymol. 409:329–345.

Suzuki, S.K., J.B. Kelley, T.C. Elston, and H.G. Dohlman. 2021. Gradient Tracking by Yeast GPCRs in a Microfluidics Chamber. In G Protein-Coupled Receptor Screening Assays. 275-287.

Wang, Y., and H.G. Dohlman. 2004. Pheromone signaling mechanisms in yeast: a prototypical sex machine. Science. 306:1508–1509.

Wang, Y., L.A. Marotti, Jr., M.J. Lee, and H.G. Dohlman. 2005. Differential regulation of G protein alpha subunit trafficking by mono- and polyubiquitination. The Journal of biological chemistry. 280:284–291.

Yu, H., P. Braun, M.A. Yildirim, I. Lemmens, K. Venkatesan, J. Sahalie, T. Hirozane-Kishikawa, F. Gebreab, N. Li, N. Simonis, T. Hao, J.F. Rual, A. Dricot, A. Vazquez, R.R. Murray, C. Simon, L. Tardivo, S. Tam, N. Svrzikapa, C. Fan, A.S. De Smet, A. Motyl, M.E. Hudson, J. Park, X. Xin, M.E. Cusick, T. Moore, C. Boone, M. Snyder, F.P. Roth, A.L. Barabasi, J. Tavernier, D.E. Hill, and M. Vidal. 2008. High-Quality Binary Protein Interaction Map of the Yeast Interactome Network. 322:104–110.

